# Natural killers in pregnancy: servants of two masters

**DOI:** 10.1101/2025.11.11.685990

**Authors:** Sergey Petrovich Krechetov, Olga Vladimirovna Khoroshkeeva, Tatjana Eduardovna Jankevic, Yuri Valentinovich Shirokov, Pavel Igorevich Borovikov, Liubov Valentinovna Krechetova, Nana Kartlosovna Tetruashvili, Dmitry Yurievich Trofimov, Gennady Tikhonovich Sukhikh

## Abstract

The absence of maternal immune rejection of a haploidentical fetus remains unexplained. We hypothesize that the presence of different HLA antigens on trophoblasts and maternal cells via killer cell immunoglobulin-like receptors (KIRs) provides two-way tuning (education) of decidual natural killer cells (dNK) with the appearance of allo- (HvG) and autoreactive (HvH) dNK. Disturbances in dNK_HvG_ and dNK_HvH_ representation may lead to abnormalities in placental development and pregnancy pathology. Our data show that recurrent pregnancy loss is not associated with specific HLA and KIR genotypes in mother and fetus, although the peculiarities of tuning involving KIR3DL2 may affect pregnancy outcome. Mathematical modeling shows the dependence of dNK_HvG_ and dNK_HvH_ representation on the probability of random KIR gene expression. The new description of NK cell involvement in the immune response in the placenta via two-way tuning is applicable to all cases where there are target cells with normal and different “guest” HLA expression (viral infections, tumors, etc.)

**Graphic abstract (Cover work):** 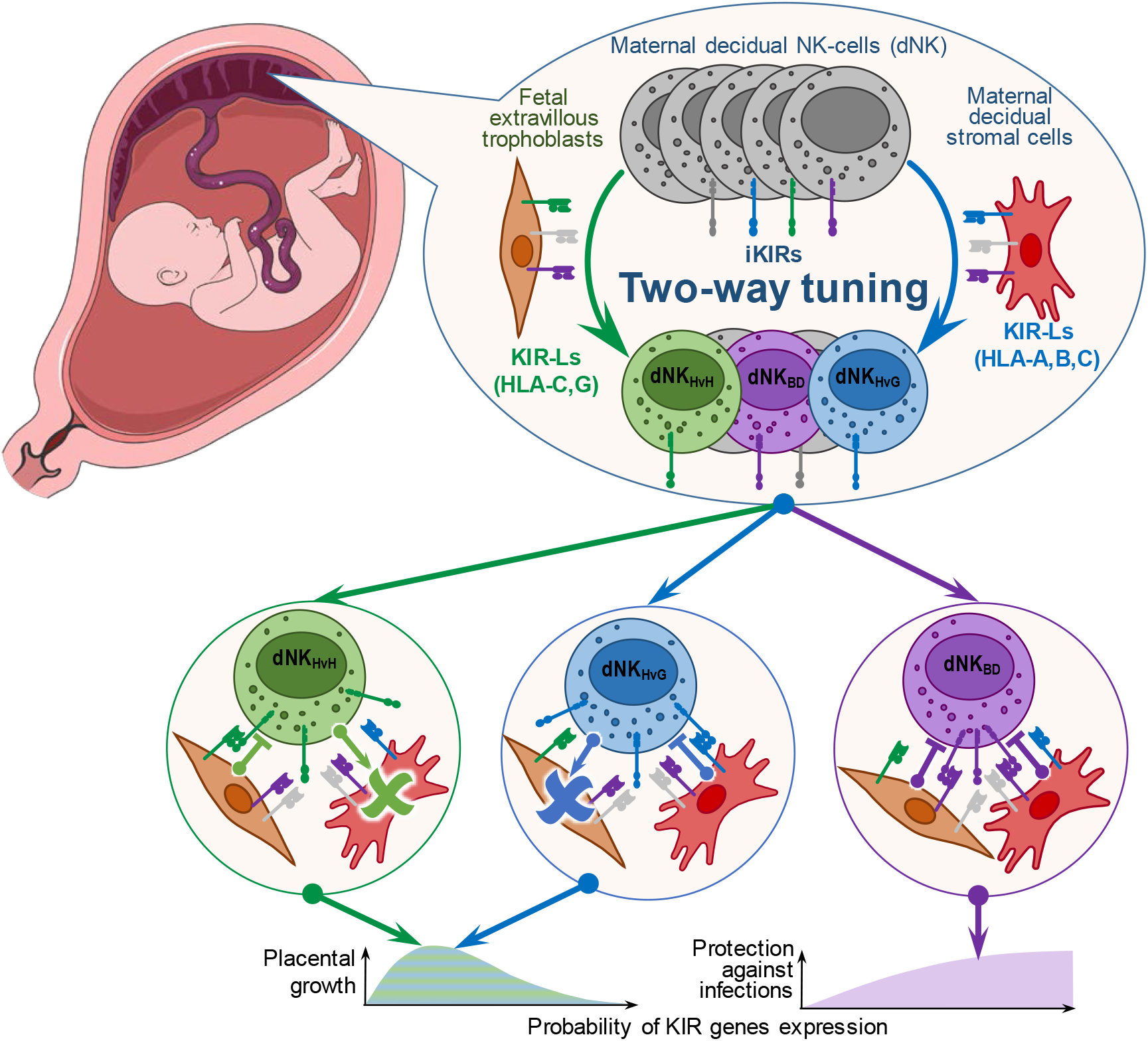

Image of fetus in utero in graphic abstract adapted from Servier Medical Art (https://smart.servier.com), licensed under CC BY 4.0 (https://creativecommons.org/licenses/by/4.0/)

**Highlights:** A hypothesis has been proposed according to which random expression of NK cell receptor genes in combination with two-way tuning (education) of NK cells by different affinity pairs of inhibitory killer cell immunoglobulin-like receptors (KIRs) and their ligands on decidual cells and trophoblasts leads to the presence of populations of maternal decidual NK cells with allo- and autoreactivity in the placenta.

It has been shown that the features of the HLA and KIR genotypes of the mother and fetus are not the underlying cause of habitual miscarriage, although genetic features can affect the course of pregnancy.

Model calculations of the representation of auto- and alloreactive maternal decidual NK cells show that the effectiveness of their participation in the structural and functional reorganization of the placenta should depend on the probability of KIR gene expression.

The hypothesis of two-way tuning of NK cells in the presence of different target cells with normal and different “guest” HLA expressions is applicable to describe the involvement of the innate immune system in the response to pregnancy, viral infection, inflammation, wounds, tumors, and transplantation.

## Main

Pregnancy is a natural phenomenon in which allorecognition does not lead to rejection but rather creates conditions for the development of a fetus immunologically foreign to the mother. Limited expression of classical human leukocyte antigen class I (HLA-I) molecules on fetal placental cells reduces their allorecognition by maternal T cells of the adaptive immune system1. However, in addition to participating in the interaction with T cell receptors, HLA-I molecules are ligands for receptors of innate immune cells - natural killer (NK cells)2, which have a cytolytic effect on cells lacking HLA-I expression. This ability of NK cells is called the “missing self” response3 and ensures the maintenance of cellular homeostasis in the body by removing cells with reduced HLA-I expression during viral infections4, oncogenic transformations5, and inflammations6. A large group of NK cell receptors interacting with classical HLA-I molecules are inhibitory and activating killer cell immunoglobulin-like receptors (iKIRs and aKIRs)7. The location of the HLA-I and KIR genes in different chromosomes (6 and 19, respectively) is combined with high allelic polymorphism of these genes8,9 and with polymorphism in the representation of KIR genes in haplotypes10. As a result, independent inheritance of the HLA-I and KIR genes leads to a wide variety of random combinations of these genes in individuals11. This feature suggests the possibility that the mother may acquire a combination of HLA-I and KIR that could cause pregnancy complications, as decidual NK cells (dNK) account for up to 70% of leukocytes in the placenta in the first trimester12. A number of studies have demonstrated a link between pregnancy complications and the presence of a combination of the AA KIR genotype in the mother and alleles of the HLA-C gene with the C2 epitope in the fetus13. It is believed that the predominance of iKIRs over aKIRs in the AA genotype, and, primarily, the presence of KIR2DL1, leads to a strong inhibitory control signal from extravillous trophoblasts (EVTs) and impaired participation of maternal decidual NK cells (dNKs) in placentation14. It should be noted that, at the same time, there are studies that do not confirm a higher rate of pregnancy failures with a combination of the inhibitory AA KIR genotype in the mother and HLA-C with the C2 epitope in the fetus15. However, the approach based on the description of NK cell control only through a simple summation of inhibitory and activating signals from KIRs16 does not explain the mechanism of identifying “missing self” in the context of the existing diversity of combinations of KIRs and KIR ligands (KIR-Ls) in individuals. A more attractive viewpoint is that the state of NK cells required for “missing self” control arises as a result of their tuning (education, licensing) through interaction with surrounding cells of the body17. The interaction of iKIRs with affine iKIR-Ls plays a decisive role in the tuning of NK cells.18. The stochastic nature of KIR genes expression results in the presence of NK cell populations with different KIRs acquisition, suggesting different tuning and involvement in “missing self” control for NK cells of different populations19. During pregnancy, dNK tuning in the placenta occurs in the presence of two types of target cells: maternal, primarily decidual stromal cells (DSCs), and fetal, primarily EVTs. Haploidentity of mother and fetus, as well as incomplete HLA-I expression on EVTs, determine differences in the KIR-L sets expressed on DSCs and EVTs (Fig.1a). This suggests that in the absence of fetal iKIR-Ls affine for maternal iKIR, dNKs of some populations may perceive EVTs as “missing self” and exhibit host-versus-graft (HvG) alloreactivity. At the same time, some dNK populations, incapable of maternal cell-mediated tuning due to the mother’s lack of affine iKIR-Ls, can be tuned by interaction with EVTs possessing these iKIR-Ls and exhibit host-versus-host (HvH) autoreactivity. This feature of dNK-mediated immune recognition in the placenta suggests the presence of two-way cytolytic activity in the placenta, which should contribute to the restriction of trophoblast expansion (HvG alloreactivity) and to endometrial remodeling (HvH autoreactivity).

**Fig. 1:**
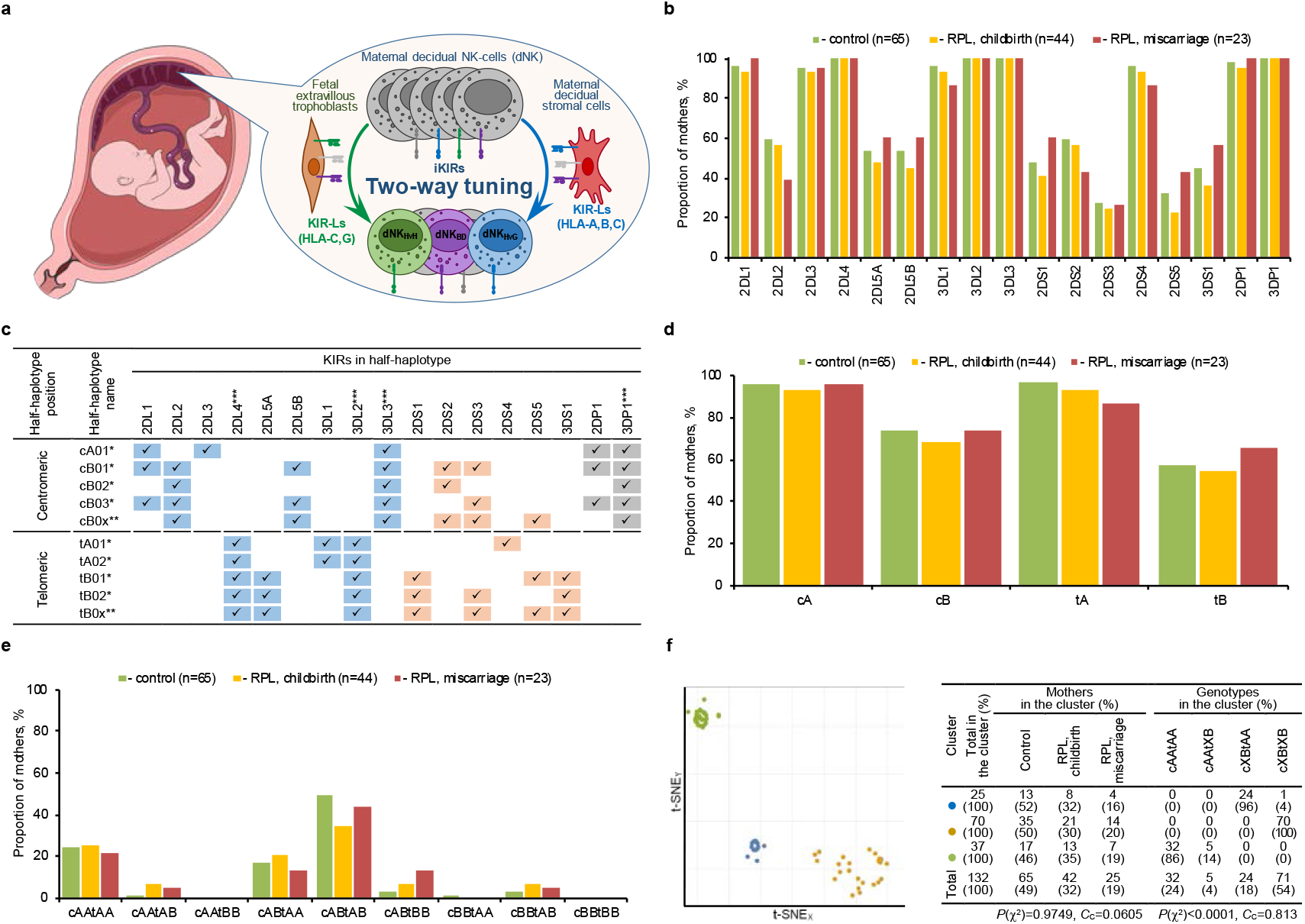
No special maternal KIR genotypes are found in cases of RPL. **a**, Schematic diagram showing the bidirectional “missing self” control by dNK in the placenta. Image of fetus in utero adapted from Servier Medical Art (https://smart.servier.com), licensed under CC BY 4.0 (https://creativecommons.org/licenses/by/4.0/). **b**, Proportion of mothers with KIR genes in clinical groups. According to the χ^2^ test (*P*(χ^2^) > 0.05), there are no statistically significant differences between the clinical groups for all genes. **c**, Composition of KIR half-haplotypes. ✓ – the gene is present in a half-haplotype. * - main half-haplotypes identified in the literature53,54,55. ** - additional half-haplotypes, which will contain at least one of the marked genes in the case that the main half-haplotypes are not identified (according to the recommendations of the Allele Frequency Net Database [http://www.allelefrequencies.net/kir6001a.asp] and IPD–the immuno polymorphism database [https://www.ebi.ac.uk/ipd/kir/about/haplotypes/]). *** - framework genes. **d**, Proportion of mothers with combinations of KIR genes included in half-haplotypes A and B according to **c. e**, Proportion of mothers with KIR genotypes determined by half-haplotypes identified in **e**. There are no statistically significant differences between the clinical groups according to the χ^2^ test (*P*(χ^2^) > 0.05) for all half-haplotypes (**d**) and all genotypes (**e**). **f**, Clustering of mothers in the space of KIR genes (left diagram) and distribution of patients and genotypes in clusters (right table). The percentage of the total number of patients in the corresponding cluster is shown in parentheses in the cells of the table. The use of XB in the designation of genotypes through half-haplotypes means the summation of genotypes with the corresponding half-haplotypes AB and BB. The results of the χ^2^ tests are shown under the table data on the clusters distribution of patients from clinical groups and patients with genotypes.

Here, using HLA-I and KIR genes typing, we demonstrate the absence of systematic genetic differences in maternal and fetal KIR and KIR-L and their combinations in pregnancies of healthy women and women with recurrent pregnancy loss (RPL). Analyzing the two-way tuning of maternal dNKs with iKIR repertoires expected due to random KIR gene expression, we find no differences between clinical pregnancy groups, but note the importance of additional HLA-G expression for ensuring HvH autoreactivity. Given the identified lack of a genetic basis for the distinct qualitative composition of KIR repertoires in pregnancies with RPL, we emphasize the importance of assessing the quantitative characteristics of the distributions of dNK functional phenotypes associated with these repertoires and dependent on the probability of KIR genes expression.

## Results

### KIR genotypes have no specific features in RPL

Genotyping results revealed no statistically significant differences in KIR gene carriage between either mothers (Fig.1b) or fetuses (Extended data Fig.1a) from different clinical groups. A comparison of clinical groups by the frequency of carriers of typical half-haplotypes (Fig.1c) and KIR genotypes, even using the extended description proposed by the authors, also did not reveal any RPL-specific features in either mothers (Fig.1d,e) or fetuses (Extended data Fig.1b,c). According to the obtained data, KIR genotypes mainly belong to mixed genotypes of different types, and the inhibitory genotype cAAtAA (coincides with AA, see Supplementary Table 2) was detected in 20-30% of patients, and its representation in the clinical groups does not have statistically significant features. Evaluation of the presence of specific KIR combinations in RPL by clustering in the KIR genes space with the allocation of 3 clusters by the number of clinical groups did not reveal (*P*(χ^2^) > 0.05) a link between the identified clusters and clinical groups in either mothers (Fig.1f) or fetuses (Extended data Fig.1d). At the same time, the linkage of clusters to KIR genotypes is highly significant (*P*(χ^2^) < 0.0001), since the presence of a certain combination of KIR genes is the basis for genotype classification.

### KIR-Ls present in the placenta have no specific features in RPL

The results of HLA-I genes typing followed by KIR-L (Fig. 2a) determination revealed no significant differences in the frequency of individual KIR-L and genotypes of HLA-C (Fig. 2b) in mothers of different clinical groups. Clustering of mothers in the KIR-Ls space (Fig. 2c) did not reveal any specific combinations of maternal KIR-Ls in RPL due to the lack of association of clusters with clinical groups (*P*(χ2) > 0.05), but showed a statistically significant (*P*(χ^2^) < 0.0001) association of these clusters with HLA-C genotypes. There is also no association (*P*(χ^2^) > 0.05) of clinical groups with clusters (Fig. 2d) formed by clustering mothers in the combined space of their KIR-Ls and KIRs, indicating the absence of specific combinations of maternal KIR-Ls and KIRs in RPL. However, statistically significant (*P*(χ^2^) < 0.0001) association of such clusters with KIR genotypes is maintained. Fetal HLA-I genes typing also revealed no statistically significant differences in the frequency of individual KIR-L and HLA-C genotypes (Fig. 2e) when restricted to HLA-I expressed by EVTs. However, when considering all KIR-Ls, a statistically significant low frequency of HLA A3/A11 not expressed on EVTs is revealed in fetuses in the RPL with miscarriage group compared to fetuses in the control group and the RPL with childbirth group (Extended data Fig. 2a). The restriction of KIR-Ls expression on EVTs to only HLA-C and HLA-G makes it unreasonable to search for specific combinations of fetal KIR-Ls in RPL using clustering in the space of fetal KIR-Ls expressed on EVTs. At the same time, this limitation does not exclude the possibility of the existence of specific combinations of maternal KIRs and fetal KIR-Ls. However, clustering in the combined space of maternal KIR and fetal KIR-Ls does not reveal any association of clusters (*P*(χ^2^) > 0.05) with clinical groups (Fig. 2f), but does retain a statistically significant (*P*(χ^2^) < 0.0001) association of clusters with maternal KIR genotypes. Assessing the presence of specific combinations of fetal KIR-Ls in RPL using clustering of fetuses in the space of all their KIR-Ls (Extended data Fig. 2b) does not reveal any association of clusters with clinical groups (*P*(χ^2^) > 0.05), but does show a statistically significant (*P*(χ^2^) < 0.0001) association of these clusters with HLA-C genotypes.

**Fig. 2:**
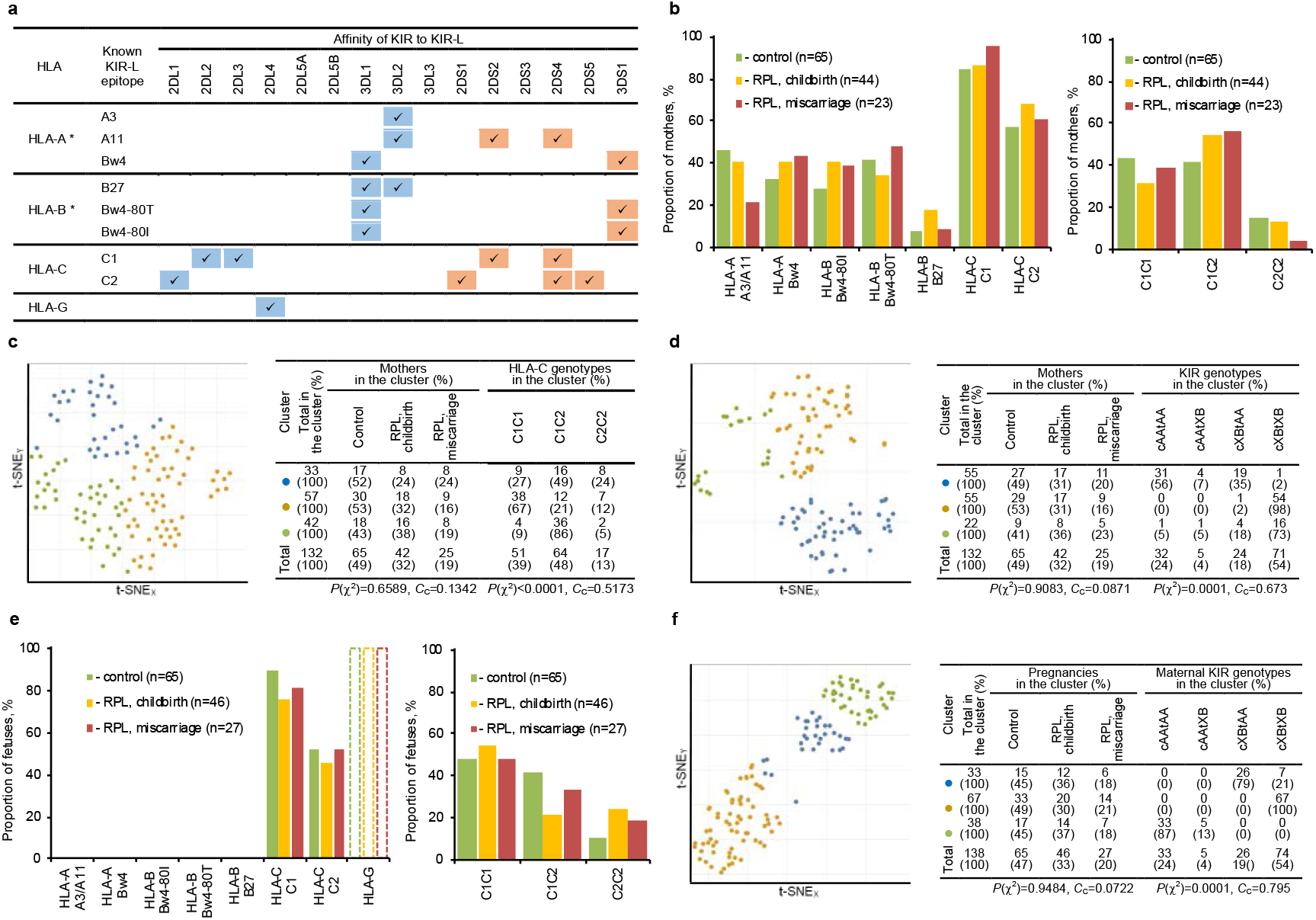
Placenta-expressed maternal and fetal KIR-Ls do not have RPL-specific features. **a**, KIR-Ls analyzed in the article56,57,58. * - for these HLA molecules there are alleles that do not contain KIR-L motifs. Cells of table for KIR and KIR-L pairs with known affinity contain the sign “✓” and are also highlighted in blue for iKIR and brown for aKIR. **b**, Proportion of mothers with KIR-Ls (left diagram) and HLA-C genotypes (right diagram). According to the χ^2^ test (*P*(χ^2^) > 0.05), there are no statistically significant differences between the clinical groups for all KIR-Ls and HLA-C genotypes. **c**, Clustering of mothers in the space of their own KIR-Ls (left diagram) and distribution of patients and HLA-C genotypes in clusters (right table). **d**, Clustering of mothers in the combined space of their own KIR-Ls and KIRs (left diagram) and distribution of patients and maternal KIR genotypes in clusters (right table). **e**, Proportion of KIR-Ls (left diagram) and HLA-C genotypes (right diagram) in fetus. Only EVT expressed KIR-Ls are shown. HLA-Gs are shown as KIR-Ls present in all fetuses due to the mandatory presence of their genes in the genotype, but the bars have a dotted border because these genes were not typed in this study. According to the χ^2^ test (*P*(χ^2^) > 0.05), there are no statistically significant differences between the clinical groups for all KIR-Ls and HLA-C genotypes. **f**, Clustering of pregnancies in the combined data space of maternal KIRs and fetal KIR-Ls expressed on EVTs (left diagram) and distribution of patients and maternal KIR genotypes in clusters (right table). On **c, d** and **f** the percentage of the total number of patients in the corresponding cluster is shown in parentheses in the cells of the table. The results of the χ^2^ tests are shown under the table data on the clusters distribution of patients from clinical groups and patients with genotypes. On **d** and **f** the use of XB in the designation of genotypes through half-haplotypes means the summation of genotypes with the corresponding half-haplotypes AB and BB.

### Genetic basis of dNK control via interactions of their KIRs with maternal and fetal KIR-Ls has minor differences in RPL

The presence of maternal KIRs, which have affine KIR-Ls of maternal and fetal origin and provide control signals for dNK from DSC and EVT, according to genotyping data, differs little between the clinical groups (Fig. 3a,b). The only statistically significant difference is the low proportion of mothers who have KIR3DL2 pairs with their own A3/A11 and/or B27 in the group of patients with RPL and miscarriage compared with the control group and the group of patients with RPL and childbirth (Fig. 3a). The main contribution to the observed difference between the groups is made by the reduced occurrence of A3/A11 in patients with RPL and miscarriage, which itself is noticeable, but statistically insignificant (Fig. 2b). Taking into account B27 led to a statistically significantly low frequency of KIR3DL2 combinations with affinity KIR-L in mothers of the RPL group with miscarriage. Estimation of the total inhibitory and total activating signals for dNK by summing all KIRs having affine KIR-L (Fig. 3c,d) reveals a statistically significant lower inhibitory signal from the fetus (Fig. 3c) if HLA-G expression is not taken into account. In addition, although both RPL groups have a lower average number of maternal iKIRs with affinity for fetal iKIR-Ls than in the control group, a statistically significant difference is observed in the pairwise comparison only for the RPL group with the childbirth. Taking into account the expression of HLA-G, a KIR2DL4 ligand, makes the differences in the overall inhibitory signal from DSCs and EVTs statistically insignificant. For maternal aKIRs in the clinical groups, a similar decrease was observed in the following order: total number, number of aKIRs with affinity to self aKIR-Ls, and number of aKIRs with affinity to fetal aKIR-Ls (Fig. 3d). However, the differences in the number of aKIRs with affine to their own aKIR-Ls and the number of aKIRs with affinity to fetal aKIR-Ls are statistically insignificant, indicating a weak effect of the absence of HLA-A and HLA-B on the total activating signal from EVTs and a similar signal level from aKIRs from both types of target cells.

**Fig. 3:**
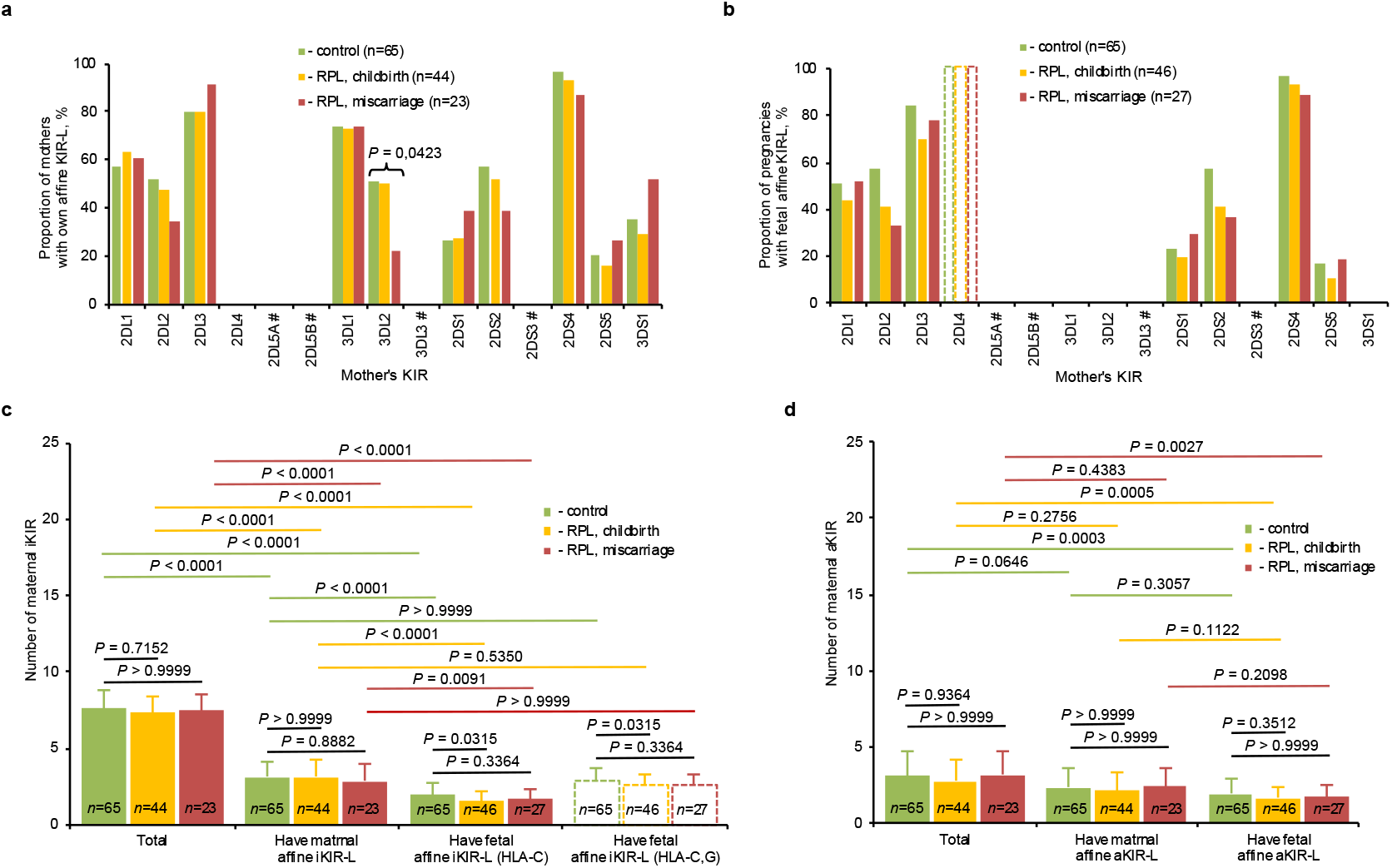
Differences in the genetic basis of two-way control of dNK via the KIR-KIR-L interaction in RPL are minor. **a**, Incidence rate of affinity pairs of KIRs and KIR-Ls in mothers (HLA-A,B,C on DSCs). Only for KIR3DL2 *P*(χ^2^) < 0.05 in comparisons of all clinical groups. Post-hoc Fisher’s exact test with Bonferroni correction show not significant differences (P > 0.017) for all pairs of clinical groups for KIR3DL2. **b**, Incidence rate of affinity pairs of maternal KIRs and fetal KIR-Ls on EVT (HLA-C,G) in pregnancies. According to the χ^2^ test, there are no statistically significant differences between the clinical groups for all KIRs (*P*(χ^2^) > 0.05). Open bars bounded by dotted lines for KIR2DL4 correspond to obligatory EVT expression of HLA-G molecules, which genes were not typed in the present study. On **a** and **b** by # are marked KIRs for which KIR-Ls are unknown. **c**, Total numbers of maternal iKIRs and number of maternal iKIRs with affine iKIR-Ls on maternal DSC (HLA-A,B,C) and fetal EVT (HLA-C,G). **d**, Total numbers of maternal aKIRs and number of maternal aKIRs with affine aKIR-Ls on maternal DSC (HLA-A,B,C) and fetal EVT (HLA-C,G). On **c** and **d** results are presented as means ± standard deviation and Kruskal-Wallis with Dunn’s post-hoc test was used for the statistical analysis.

### In some pregnancies, maternal and fetal genotypes suggest the absence of dNKs cytolytic for EVTs, and this is more often found in RPL

Analysis of the presence and number of combinations of iKIRs with different functional phenotypes (Fig. 4a,b,c,d) in pregnancies, in the case of iKIR-Ls restriction to HLA-A, B and C (HLA-ABC) on DSCs and HLA-C on EVTs, reveals a statistically significant trend towards a more frequent absence of dNK_HvG_ in RPL (Fig. 4e). Pregnancies negative for dNK_HvG_ were detected in all clinical groups, but their number was higher (approximately 25%) in the RPL group with miscarriage. In the control group and the RPL group with the childbirth, pregnancies lacking dNK_HvG_ occurred in a similar proportion of cases and did not exceed 10%. Restricting iKIR-Ls expression in EVTs to HLA-C suggests a low proportion of pregnancies (less than 20%) with dNK_HvH_. Taking into account the presence of HLA-G in all EVTs (Fig. 4f) does not change the estimate of the proportion of pregnancies with the presence of dNK_HvG_, but leads to 100% of pregnancies with the presence of dNK_HvH_. The proportions of iKIR gene combinations for each of the functional phenotypes dNK_hypo_, dNK_HvG_, and dNK_HvH_ are significantly lower than the proportion of combinations for the dNK_BD_ phenotype for all pregnancies and do not differ between clinical groups (Fig. 4g,h). When dNK tuned by only HLA-C expression on EVTs (Fig. 4g) the average proportion of dNK_HvH_ combinations does not exceed a few percent and is statistically significantly lower than the proportions of other phenotypes in all clinical groups. With the addition of HLA-G expression, the proportion of dNK_HvH_ combinations increases up to 10% due to some dNK_hypo_ combinations, which aligns it with the proportions of combinations for dNK_hypo_ and dNK_HvG_ (Fig. 4h). Moreover, the proportion of dNK_HvG_ combinations when taking into account HLA-G expression decreases due to the classification of some combinations as BD. In general, HLA-ABC on DSCs combined with HLA-C and HLA-G on EVTs during pregnancy ensures 100% presence of dNK_hypo_, dNK_HvH_, and dNK_BD_ in the placenta. However, pregnancies in which dNK_HvG_ are absent are possible.

**Fig. 4:**
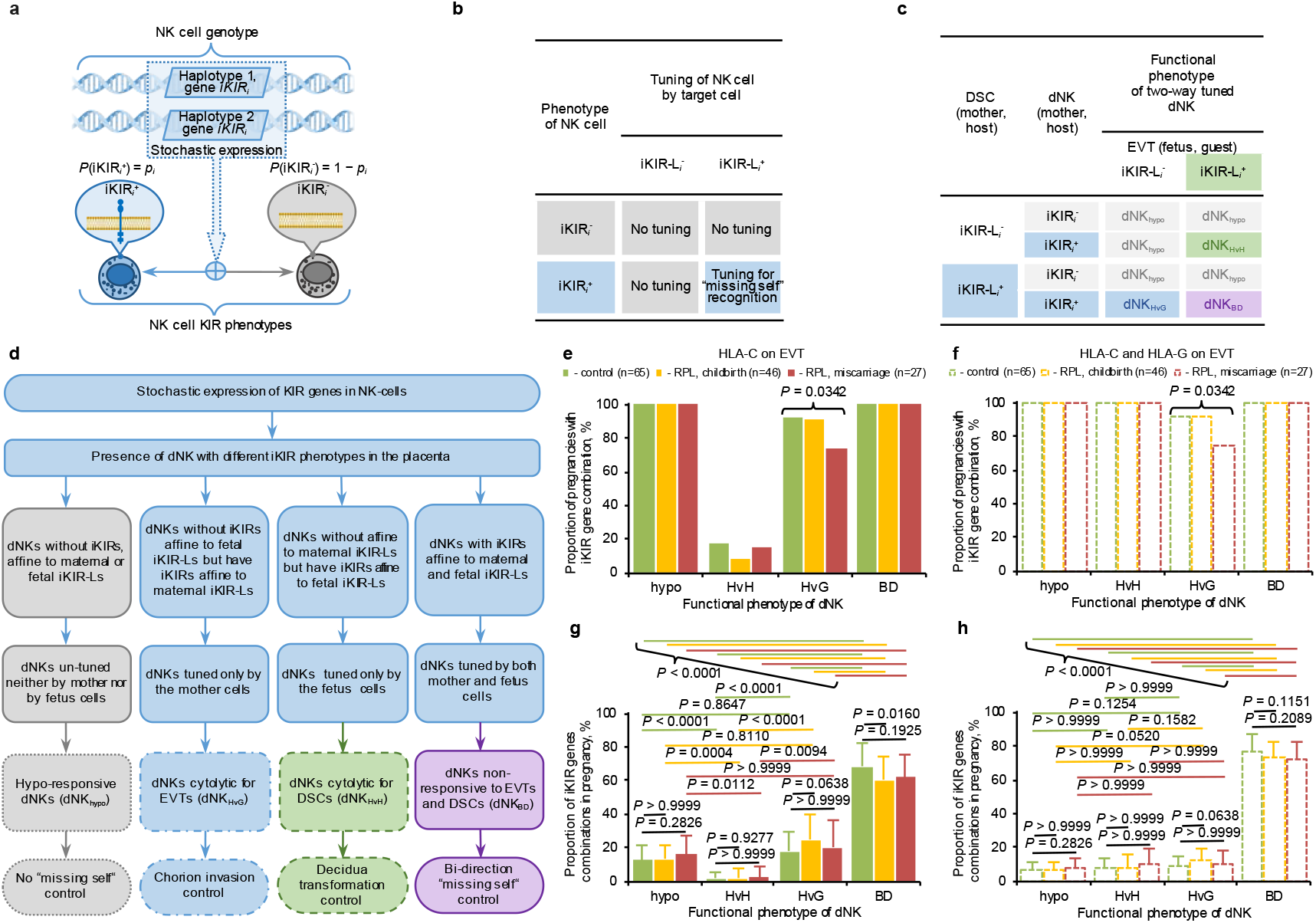
In RPL, there is minor genetic basis for abnormal representation of dNK functional phenotypes that can be generated by two-way tuning (education). **a**, Schematic description of the emergence of various iKIR phenotypes of NK cells as a result of stochastic expression of KIR genes. Phenotypes with iKIR_*i*_ acquisition (iKIR_*i*_^+^) and iKIR_*i*_ absence (iKIR_*i*_^-^) are shown. The acquisition of iKIR_*i*_ occurs with probability *p*_*i*_ due to stochastic expression of the *iKIR*_*i*_ gene (NIAID Visual & Medical Arts. (10/7/2024). DNA. NIAID NIH BIOART Source. bioart.niaid.nih.gov/bioart/123). **b**, NK cell tuning (education) mediated by the interaction of its iKIR_*i*_ and iKIR-L_*i*_ of target cell. iKIR-L_*i*_^+^ - HLA allele with iKIR_*i*_ affine epitope, iKIR-L_*i*_^-^ - HLA allele without iKIR_*i*_ affine epitope. Tuning take place only if immunological synapse is formed by NK cell that is iKIR_*i*_^+^ and target cell that is iKIR-L_*i*_^+^. **c**, The two-way tuning of the dNK carrying iKIR_*i*_ gene in chimeric microenvironment of target cells including DSC (mother, host) and EVT (fetus, guest). Gray dNK_hypo_ – un-tuned hyporeactive dNK; green dNK_HvH_ – tuned dNK, cytolytic against DSCs and “missing self” EVTs; blue dNK_HvG_ – tuned dNK, cytolytic against EVTs and “missing self” DSCs; purple dNK_BD_ - tuned dNK, cytolytic against “missing self” DSCs and EVTs (bidirectional control of “missing self”). **d**, Hypothetical role of dNK with different functional phenotypes in the placenta. **e - f**, Proportion of pregnancies with at least one iKIR gene combination for dNK functional phenotypes when dNK tuning by EVT is restricted to HLA-C (**e**) or involves HLA-C and HLA-G (**f**). On **e** and **f** only for HvG phenotype *P*(χ_2_) < 0.05 in comparisons of all clinical groups. Post-hoc Fisher’s exact test with Bonferroni correction show not significant differences (P > 0.017) for all pairs of clinical groups for HvG. **g - h**, Proportions of iKIR genes combinations for dNK functional phenotypes in pregnancies when dNK tuning by EVT is restricted to HLA-C (**g**) or involves HLA-C and HLA-G (**h**). On **g** and **h** results are presented as mean ± standard deviation. *P* values were determined using Kruskal–Wallis tests followed by Dunn’s post hoc test for multiple comparison of clinical groups and using Friedman test with Dunn’s post hoc test for multiple comparison of functional phenotypes.

### The absence of a clear link between RPL and genetics of HLA and KIR indicates the rationale for searching causes of this pregnancy pathology in the control of KIR expression

Obviously, information on maternal and fetal HLA and KIR genes only allows us to determine the presence and number of KIR gene combinations whose expression implies the presence of dNK_hypo_, dNK_HvG_, dNK_HvH_, or dNK_BD_ (Fig. 4e-h). However, the proportion of dNKs with a particular functional phenotype depends on the parameters of random iKIR genes expression, which determines the probability of dNK acquiring each possible iKIR combination. In the context of an approximate estimation, it is advisable to assume that the expression of iKIR genes is independent and identical, characterized by the probability *p* of expression of each gene (See Methods for details). Fig. 5a and Supplementary Fig. 3 show examples of the dependences on *p* of the expected values *E*_F_ (*p*) of the proportions of dNKs with different functional phenotypes (*E*_hypo_ (*p*), *E*_HvG_ (*p*), *E*_HvH_ (*p*) и *E*_BD_ (*p*)) for individual pregnancies with different *N*_iKIR_ and different combinations of iKIR-L on DSCs and EVTs. The type of dependences on *p* of the average expected value *E*_F_ (*p*), calculated for all pregnancies in the clinical groups (Fig. 5b), reflects the fact that the function *E*_hypo_ (*p*) is monotonically decreasing, the function *E*_BD_ (*p*) is monotonically increasing, and the functions *E*_HvG_ (*p*) и *E*_HvH_ (*p*) have a maximum at *p* no more than 0.5. The indicated types of functions *E*_F_ (*p*) are due to the fact (Supplementary Fig. 4a,c,e) that the functional phenotypes HvG or HvH can predominantly acquire dNK with one or two iKIRs, the phenotype hypo can mainly be possessed by dNK without iKIRs or with one iKIR, and the phenotype BD cannot occur in the absence of iKIRs and predominates when dNK acquires two or more iKIRs. Since with an increase in *p*, a shift to the right of the probability function of the binomial distribution of the number of receptors acquired by dNK occurs (Supplementary Fig. 4b,d,f), and the resulting type of functions *E*_F_ (*p*) arises. Comparison of clinical groups by the expected values of dNK functional phenotypes acquisition at different *p* values (Fig. 5b) revealed no statistically significant differences between the groups. This again points to, if specific features of dNK alloreactivity play a role in the development of RPL, this is poorly related to specific HLA and KIR genotypes. Mathematical modeling of the influence of random KIR genes expression in dNKs on the enrichment by functional phenotypes points to distinctions of maternal KIR genes expression as a possible cause of altered dNK alloreactivity in RPL (Fig. 5c).

**Fig. 5:**
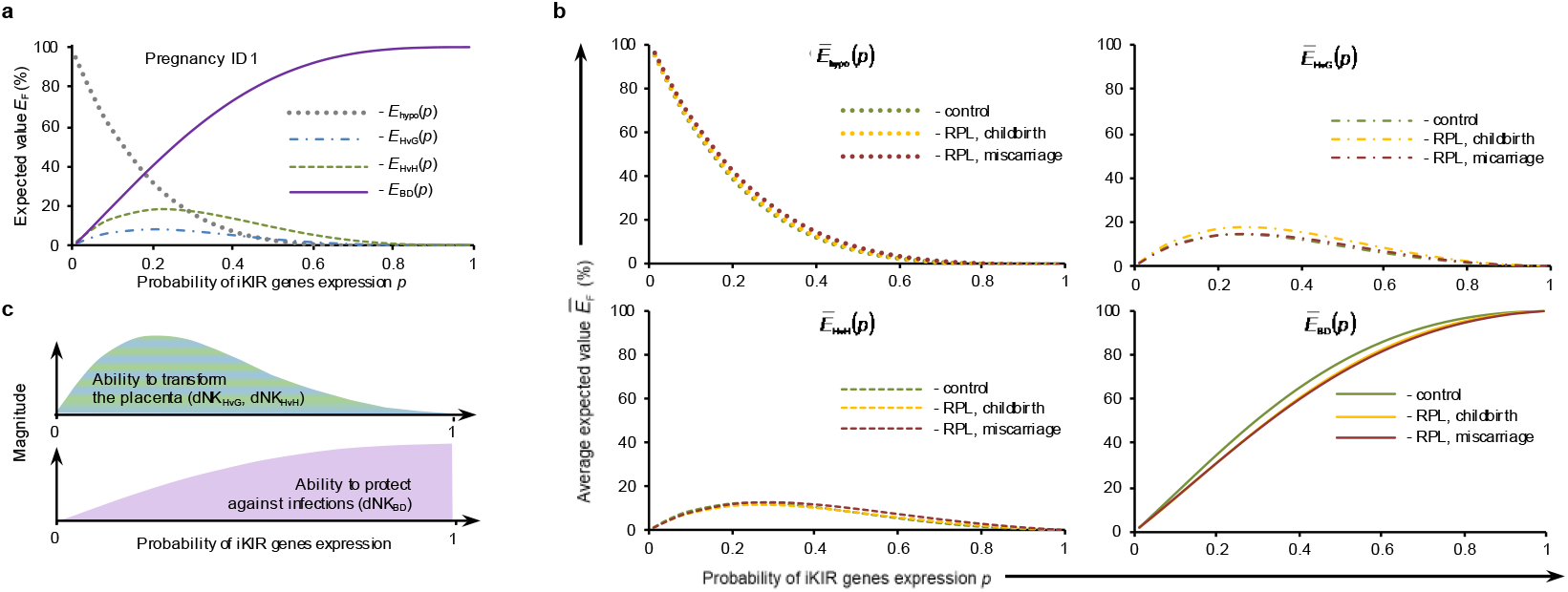
Not genetic features, but under- or overexpression of KIR genes may be the cause of placental disorders in RPL. **a**, Hypothetical expected value *E*_F_ (*p*) of the proportions of dNK functional phenotypes at different probabilities *p* of iKIR genes expression for individual physiological pregnancy #1. Detailed information about this example pregnancy #1 is given in Supplementary Fig.3a. **b**, Average expected values 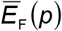 of the proportions of dNK functional phenotypes at different probabilities of iKIR genes expression in clinical groups (65 cases for control; 46 cases for RPL and childbirth, 27 cases for RPL and miscarriage). In comparing clinical groups, Kruskal-Wallis test reveals no significant differences (*P* > 0.05) for *E*_F_ (*p*) of all functional phenotypes at *p* = 0.01, 0.1, 0.2, 0.3, 0.4, 0.5, 0.6, 0.7, 0.8, 0.9, 0.99. Calculations for **a** and **b** were made assuming expression of HLA-A,B,C on DSC and HLA-C,G on EVT. See Methods for *E*_F_ (*p*) and 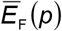 calculation details. **c**, Hypothetical influence of iKIR genes expression probability on participation of dNK in the transformation of the placenta (green-blue gradient fill) and elimination of infected and damaged “missing self” cells (purple fill).

## Discussion

The data obtained in this and other researches show that HLA and KIR genotypes are not the main cause of RPL, although they may influence pregnancy. The lack of a link between the cause of RPL and the genetics of the main allogenic molecules makes it advisable to search for the cause in the mechanism of inclusion of HLA and KIR molecules in the realization of immune functions. To perform surveillance functions in both adaptive (T and B cells) and innate (NK cells) immunity, naive immunocompetent cells, with or without the necessary properties, are randomly generated in the body. The existence of NK cells in different functional states is associated with the random (stochastic) acquisition by these cells of a set of KIRs and CD94/NKG2 receptors (killer cell lectin-like receptors)20, and possibly, LILRs (leukocyte Ig-like receptors)21. Random expression of KIR genes is provided by special promoters acting as probabilistic switches22. The importance of the mechanism of random expression of receptors playing a role in the tuning of NK cells is indicated by the presence of this feature in receptors of the Ly29 family23, which have the same role in mouse NK cells as KIR in human24. Moreover, both the structure of the Ly29 and KIR molecules and the structure of their gene promoters are different25. There are no data on the functioning of the CD94/NKG2 and LILR gene promoters as probabilistic switches26, however, the CD94/NKG2 and LILR receptors, like KIR, are not expressed on all NK cells and are randomly combined with each other27.

Random KIR expression exhibits features of a dependent nature of receptor co-expression, which is manifested in the deviation of experimental data from calculations of the probability of the presence of several receptors in the set acquired by an NK cell according to the “product rule”28. This feature of iKIRs can be explained by the existence of a specific mechanism controlling the co-expression of closely located genes29. However, allowing for variation in the probability *p* of KIR expression between NK cells makes it possible to explain the observed deviation from the “product rule” without invoking a special mechanism for controlling co-expression. In particular, if in the binomial distribution *X ∼ B*(*N*_iKIR_, *p*) *p* is randomly drawn from a beta distribution *p ∼ Beta*(*α, β*), this leads to the joint beta-binomial distribution *X ∼ BetaBin*(*N*_iKIR_,*α, β*) 30, which is proposed as the most appropriate to describe the distribution of the amount of KIR acquired by NK cells28. It is interesting to note that the different shapes of the probability density function for the beta distribution (Supplementary Fig. 5a,b,c) assume different magnitudes of differences between *P*(*X* = *n*) according to the beta-binomial distribution from *P*(*X* = *n*) according to the product rule based on the binomial distribution (Supplementary Fig. 5d,e,f). In addition, for the same average values of the probability of expression of individual genes, a different type of distribution of this probability should correspond to a different representation of functional phenotypes of dNK during pregnancy (Supplementary Fig. 5g,h,i). The identification of individuals with different probabilities of KIR expression by NK cells28 indicates a significant variability of this parameter in humans and the necessity of its study in RPL. The data on the decrease in KIR expression on dNK during pregnancy31, reflecting a decrease in the expression probability, in light of the discussed random acquisition of an KIR set can be interpreted as a shift to a state with the expression of a limited number of iKIRs on an individual dNK (Supplementary Fig. 4b,d,f) and a higher representation of dNK_HvH_ and dNK_HvG_ among tuned dNK (Fig. 5a,b). However, according to modeling, the dependence of the proportions of dNK_HvH_ and dNK_HvG_ in the placenta on the probability of KIR expression has a maximum (Fig. 5a,b), which allows for different effect of an additional decrease in this probability during pregnancy. As a result, an additional reduction in KIR expression in RPL and preeclampsia32,33 may lead to cytolytic dNK activity that is unconformable for the remodeling of placental tissue and blood vessels to meet the needs of the growing fetus.

The proposed description of NK cell involvement in alloimmune interactions with two-way tuning in the placenta is well applicable to the description of the immune response in virus-infected tissue. A distinctive feature of virus-infected target cells is the increased expression of HLA-G with suppressed expression of HLA-ABC, which distinguishes them from healthy cells with full expression of only HLA-ABC34. The emerging difference in the iKIR-L sets in healthy and infected cells suggests the existence of a two-way tuning of NK cells in the infected tissue and, if we conditionally call the infected target cells “guests”, then in the infected tissue we can conditionally distinguish NK_hypo_, NK_HvG_, NK_HvH_ and NK_BD_ by analogy with the approach proposed for dNK (Fig. 4c). It should be noted that it is possible to talk about the participation of the KIR2DL4 and HLA-G pair in the tuning of NK cells only conditionally. We are unaware of direct experimental data on the tuning in NK cells expressing KIR2DL4 under conditions of HLA-G expression on target cells. However, the ability of HLA-G on target cells to inhibit their cytotoxic damage by NK cells35 allows to assume the involvement of the KIR2DL4 and HLA-G pair in tuning and to speak of the emergence of NK_HvH_ as a result of tuning of NK cells expressing KIR2DL4. The assumption of participation in the tuning of a KIR2DL4, which has inhibitory properties36, suggests that two-way tuning will be a mandatory feature of the “guest” expression of HLA-G due to the framework *KIR2DL4* gene. According to a common view, the observed changes in HLA-I expression on infected cells lead these cells from destruction and promote viral survival37. Just as importantly, however, maintaining the viability of virus-infected cells may serve the survival of the host, as excess death of infected cells may result in significant structural and functional abnormalities in host tissues and organs. In this regard, “guest” expression should be considered a protective mechanism preventing excessive activity of killer T cells and NK cells, which can lead to a cytokine storm with irreversible tissue damage and even death of the virus-infected host. In addition, the acquisition of the guest phenotype by host cells infected with the virus is not only a way to control the response to viral multiplication in the acute period through immunosuppression. During recovery, it is important to create conditions for the restoration of the tissue damaged by the viral load. It is necessary not only to remove cells suspicious of infection. At the same time, it is necessary to stimulate the proliferative activity of uninfected cells and remove senescent cells that interfere with renewal. In this regard, the existence of NK_HvH_ implies the acceleration of the removal of senescent cells with less resistance to damage38. Equally important is the effect of activated NK cells on the repair of damaged tissue through the inhibitory effect of IFN-γ on collagen synthesis by fibroblasts39 and through the slowing of wound healing40. This means that the presence of activated NK cells through slowing regeneration and preventing fibrosis should contribute to the prolonged restructuring of the regenerated tissue and the construction of its structure that meets the needs of the surrounding formations and the body as a whole41. In this regard, the ability of IFN-γ to induce the expression of HLA-G in cells of different origin42 and therefore maintain conditions for two-way tuning in tissues with increased NK cell activity is noteworthy.

Consideration of the “guest” expression of HLA-G as a condition for two-way tuning promoting the intensification of tissue renewal and remodeling suggests that low HLA-G expression may be the cause of insufficient reorganization of the bloodstream and limited blood supply in the endometrium, which are considered the leading factor of placental insufficiency in RPL. This assumption is consistent with the increased expression of HLA-G on trophoblasts by progesterone43 and dexamethasone44, which are effective in the treatment of RPL. It is interesting to note that mesenchymal stem cells (MSCs), considered mandatory participants in tissue healing and renewal and present in the placenta, can contribute to creating conditions for two-way tuning through the HLA-G expression45. In particular, human MSCs show an increase in HLA G expression not only in the presence of progesterone46 and dexamethasone47, but also in the conditions of hypoxia48, which accompanies the insufficiency of blood supply in the placenta. Unlike the cases discussed above with a beneficial effect on the body, “guest” HLA-G expression with the loss of classical HLA-I in tumors should be considered pathogenic. Two-way tuning in this case will lead to the involvement of NK cells in the restructuring of tumor and surrounding tissues, promoting vascularization and tumor invasion.

A special case of conditions for two-way tuning of NK cells is organ or tissue transplantation, in which there is a coexistence of NK cells and target cells with the KIR and HLA genotypes of the recipient and donor in the recipient’s body. In this case, one can formally talk about two-way tuning for NK cells of both the donor and the recipient. However, only the possible role of donor NK cell tuning to guest-versus-host (GvH) activity in the treatment of leukemia using hematopoietic stem cell transplantation is being investigated49, since the presence of NK_GvH_ may underlie the graft-versus-leukemia (GvL) reaction50. In connection with the results of our studies, it is interesting to draw attention to the data on the better renal graft engraftment in the absence of HLA-A3/A11 in the donor51. According to the model of two-way tuning, the absence of HLA-A3/A11 in the donor corresponds to a higher proportion of NK_HvG_ and/or a lower proportion of NK_HvH_ in the recipient compared to the recipient of a kidney from an HLA-A3/A11-negative donor (Supplementary Note. 1). Such a conclusion about the possible positive effect on the viability of the “guest” of a higher KIR3DL2-related alloreactivity of the HvG type is consistent with the result obtained in our study (Fig. 3a, Fig. 4e). Understanding the specific features of two-way tuning of NK cells in the donor-recipient setting is of interest for the treatment of RPL using lymphocyte immunotherapy (LIT), which has demonstrated clinical utility. The observed decrease in the number and cytolytic activity of NK cells in the maternal blood indicates the involvement of these cells in response to LIT and other types of immunotherapy52. Moreover, a decrease in the total cytolytic activity of NK cells may indirectly indicate a decrease in their KIR expression28, which corresponds to changes in dNK upon the onset of pregnancy31,32 and may be considered as an indication that LIT contributes to achieving the required decrease in KIR expression. However, the isolation of NK cells from the blood and the absence of KIR determination in the corresponding studies52 make the extension of the statement about the decrease in KIR expression under the influence of LIT to dNK ambiguous. It should be noted that, as for RPL, the clinical efficacy of LIT cannot be associated with KIR genotypes and KIR-L representation of mothers and fathers53. This once again emphasizes the importance of studying random KIR expression as a possible basis for controlling NK cell activity through changes in the representation of carriers of different KIR sets, which, after tuning, determine different abilities for a cytolytic response based on the presence of KIR-L on target cells in the microenvironment.

Overall, based on our data and analysis, we have substantiated the hypothesis that random expression of NK cell receptor genes, combined with the tuning (education) of NK cells by affine iKIR and iKIR-L pairs under conditions of existing KIR and HLA I polymorphism and polygenicity, leads to the presence of maternal dNK populations with alloreactivity and autoreactivity in the placenta. Alloreactive dNK cells inhibit trophoblast invasion, while autoreactive dNK cells promote the destruction of existing endometrial structure and its remodeling during decidualization and placental growth. However, the data obtained and the assessments made on their basis show that the presence of affinity pairs iKIR iKIR-L, as well as the presence of dNK functional phenotypes, are not specific to RPL. At the same time, according to model calculations, the representation of auto- and alloreactive dNK populations and, consequently, the effectiveness of their participation in placental structural and functional remodeling depends on the probability of KIR gene expression. The latter indicates the need for further targeted studies of the role of the characteristics of random expression of NK cell receptors, primarily KIR, in the pathogenesis of RPL.

## Methods

### Patients

The study was approved by the Local Ethics Committee of the National Medical Research Center for Obstetrics, Gynecology, and Perinatology, named after Academician V.I. Kulakov of the Ministry of Health of the Russian Federation (Moscow, RF), as complying with the current requirements of the guidelines of the Russian Federation and the World Medical Association (in accordance with the Declaration of Helsinki) (LEC of the National Medical Research Center for Obstetrics, Gynecology, and Perinatology, Decision No. 35 of April 9, 2009).

The study included 65 healthy mothers (control) and 67 women diagnosed with RPL, observed at the National Medical Research Center for Obstetrics, Gynecology, and Perinatology, named after Academician V.I. Kulakov in 2009-2024. Women enrolled in the study were aged 20 to 45 years. Control patients had one successful pregnancy prior to study entry. Patients with RPL were characterized by two or more consecutive spontaneous abortions (up to 22 weeks) from the same partner60. Exclusion criteria from the study were the presence of karyotype features in parents; anatomical causes of habitual miscarriage; chronic infectious, inflammatory, malignant oncological and systemic autoimmune diseases in mothers; maternal obesity (BMI ≥ 30); fetal malformations and/or congenital malformations in the ultrasound examination; Rh sensitization in pregnancy. Clinical history of patients enrolled in the study is summarized in Supplementary Table 1. All RPL patients received standard treatment (https://www.eshre.eu/Guidelines-and-Legal/Guidelines/Recurrent-pregnancy-loss).

All patients became pregnant naturally. According to the outcome of pregnancy, patients with RPL were divided into two groups: 44 patients who gave birth (RPL, childbirth) and 23 patients who miscarried (RPL, miscarriage). Mothers with RPL and several pregnancies during the study were assigned to the RPL with childbirth group, since everyone had at least one pregnancy with the birth of a child. Repeated pregnancies were counted as individual clinical cases, resulting in 46 pregnancies in the RPL group with a carried pregnancy and 27 pregnancies in the RPL group with a miscarriage.

### Samples

Whole peripheral blood was used as biological material in mothers and newborns for analysis. 5 ml of blood was collected from the cubital veins of mothers in Vacuette tubes (Greiner Bio-One, Kremsmünster, Austria) containing EDTA as an anticoagulant. From newborns, 3-5 ml of umbilical cord blood, flowing freely from the umbilical cord after omphalotomy, was collected in a test tube containing the same anticoagulant. In cases of miscarriage, villous chorion tissue was used as fetal biological material. After curettage of the uterine cavity, several chorionic villi with a total weight of 2-3 milligrams were placed in 0.5 ml of physiological saline and treated with proteinase K using PREP-PK kit P-028-N/2EU (DNA-Technology Research & Production LLC, Moscow, Russia). Genomic DNA was isolated from blood and chorion samples using PREP-MB MAX DNA Extraction Kit P-103-A/8EU (DNA-Technology Research & Production LLC, Moscow, Russia).

### HLA class I (HLA-I) genotyping and KIR-L determination

DNA libraries for typing the HLA-A, B, and C genes (HLA ABC) were prepared using HLA-Expert kit S6-H010-NI N v3 (NPO DNA-Technology LLC, Moscow, Russia). Sample amplification and library preparation were performed using DTprime amplifier (DNA-Technology Research & Production LLC, Moscow, Russia). The resulting library was sequenced on a MiSeq instrument (Illumina, Inc., San Diego, CA, USA) using the MiSeq Reagent Kit v3 (Illumina, Inc., San Diego, CA, USA). Based on the sequencing results, HLA I genotyping was performed using HLA-Expert version 2.0 software (DNA-Technology Research & Production LLC, Moscow, Russia). KIR-L epitopes (A3/11, Bw4, B27, and C1/C2) were determined based on the identified HLA-I alleles (https://hla.alleles.org/antigens/bw46.html, https://www.ebi.ac.uk/ipd/kir/matching/ligand/). The results of KIR-L epitope determination for all mothers and fetuses are presented in Supplementary Table 2.

### KIR genotyping and determination of KIR genotypes

The primers were designed and synthesized by “DNA-Technology Research & Production”, LLC. Using developed primers, DNA regions containing exons 3, 4, and 5 of the KIR genes and pseudogenes were amplified. Ten primers were used, including four forward and six revers primers, resulting in amplification products of 377, 354, and 449 bp in length. Sequencing of the prepared libraries was performed as described above for HLA-I gene typing. Genotyping of KIR based on sequencing results was performed using software developed from HLA-Expert version 2.0 (DNA-Technology Research & Production LLC, Moscow, Russia). The resulting sequences were compared with the reference exon sequences from the IPD-KIR database version 2.13 (https://www.ebi.ac.uk/ipd/kir/release/v213/). As a result, each mother and each fetus were assigned a binary KIR genotype (https://www.ebi.ac.uk/ipd/kir/about/nomenclature/genotype/) indicating the presence of the genes *KIR2DL1, KIR2DL2, KIR2DL3, KIR2DL4, KIR2DL5A, KIR2DL5B, KIR2DS1, KIR2DS2, KIR2DS3, KIR2DS4, KIR2DS5, KIR3DL1, KIR3DL2, KIR3DL3, KIR3DS1, KIR2DP1* и *KIR3DP* (Supplementary Table 2).

The data obtained make it possible to classify the genotypes of all patients as AA and Bx in accordance with the recommendations of the KIR Nomenclature Committee (https://www.allelefrequencies.net/kir6001a.asp). According to these recommendations, if any of the genes *KIR2DL2, KIR2DL5, KIR3DS1, KIR2DS1, KIR2DS2, KIR2DS3*, or *KIR2DS5* is present in a genotype, the genotype is considered to contain haplotype B. In this case, the AB and BB genotypes are considered indistinguishable and are designated Bx. If none of the above genes are present, the genotype is considered AA. However, the KIR Nomenclature Committee’s recommendations only highlight the potential biological and medical significance of the differences between haplotypes A and B. At the same time, they exclude a more detailed consideration of possible genotypes characteristics that may distinguish patients with RPL. We additionally classified genotypes based on the presence of centromeric and telomeric half-haplotypes A and B61. The presence of obligatory genes of typical half-haplotypes (Fig.1c) was used as an indicator of the presence of half-haplotypes A or B in a patient. The presence of half-haplotypes was determined using the algorithm given in the Supplementary Fig.1. Based on the mandatory presence of two centromeric and two telomeric half-haplotypes in the genotype, if evidence of only one type of centromeric or telomeric half-haplotype (A or B) was present, both centromeric or telomeric half-haplotypes in the genotype were considered to have this type. According to the resulting set of half-haplotypes, all genotypes were divided into eight groups based on the presence of half-haplotypes A and B: cAAtAA, cAAtAB, cAAtBB, cABtAA, cABtAB, cABtBB, cBBtAA, cBBtAB, and cBBtBB. Supplementary Table 2 presents the results of genotype determination using the authors’ algorithm and in accordance with the recommendations of the KIR Nomenclature Committee. It is noteworthy that the cAAtAA genotypes coincided with AA, and the genotypes containing at least one half-haplotype B coincided with Bx.

### Description of alloimmune response involving dNK

An algorithm for describing the expected alloimmune response involving dNK is presented in the Supplementary Fig.2. For each pregnancy, maternal iKIRs and aKIRs, for which affine KIR-Ls are present in mother and/or fetus, were determined. The hypothetic involvement of individual iKIR in dNK tuning (acquisition of dNK capacity for cytolytic response) was assessed using rules (Fig.4a,b,c), reminiscent of those used to assess donor NK cell alloreactivity during hematopoietic stem cell transplantation, based on the presence of iKIR-L in the donor and recipient62,63. It was considered that in the presence of affine iKIR-L on DSCs and its absence on EVTs, iKIR tunes dNKs to react to EVTs as “missing self” suggesting a cytolytic effect of dNKs against EVTs (host-versus-graft (HvG) response). In the presence of affine iKIR-L on EVTs and its absence on DSCs, iKIR causes tuning with the acquisition of dNK capacity for cytolytic action on maternal cells (host-versus-host (HvH) response). The presence of affine iKIR-L on both DSCs and EVTs means that iKIR will tune dNKs when interacting with both DSCs and EVTs, and dNKs will acquire the ability to protect against “missing self” cells of both types (BD): DSCs and EVTs. The absence of iKIR in dNK and/or the absence of iKIR-L on both DSCs and EVTs implies the inability to tune dNK, suggesting the existence of dNK in a hypo-responsive state (hypo) with the inability to cytolytic response to “missing self”.

Further, based on estimates of participation in the tuning of individual iKIRs, the functional phenotype of each possible combination of iKIRs, which can acquire dNK as a result of random (stochastic) expression of KIR genes, was determined. According to a combinatorial analysis, for dNK cell with a total number *N*_iKIR_ of iKIR genes there are 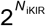 of combinations with number of iKIRs from zero to *N*_iKIR_. For each iKIR combination, the resulting dNK functional phenotype was assigned based on the premise that the presence of at least one iKIR with an affine iKIR-L on the synapse-forming target cell is sufficient to suppress cytolytic activity and tune the NK cell. At the same time, possible differences in the cytolytic potential of the response to “missing self” in the presence of different numbers of iKIR-iKIR-L affine pairs are not taken into account, and all iKIRs are considered equivalent in the effectiveness of inhibitory action and tuning. In this case, the presence of at least one iKIR with the BD status in an iKIR combination gives rise to the same functional phenotype BD in dNK (dNK_BD_), regardless of the status of the other iKIRs. The BD functional phenotype also arises in combinations that do not contain iKIRs with the BD status but include at least one iKIR with the HvG status and at least one iKIR with the HvH status. The presence in combination of one or more iKIRs with HvG status and the rest iKIRs with hypo status leads to the acquisition by dNK of the functional HvG phenotype (dNK_HvG_). One or more iKIRs with HvH status with the remaining iKIRs with hypo status in combination leads to dNK acquisition of the functional HvH phenotype (dNK_HvH_). The absence in combination of iKIR with BD, HvG or HvH status makes dNK hyporeactive (dNK_hypo_) in the “missing self” response. Using the described approach, each pregnancy was characterized by the presence and proportion of combinations of maternal iKIR genes generating a certain functional phenotype from the total number of possible combinations of detected maternal iKIR genes. Involvement of the maternal A3, A11, B27, Bw4, C1, and C2 epitopes and involvement of the fetal C1 and C2 epitopes alone or in combination with the HLA-G epitope was considered. The possible inability of KIR3DL2-single positive dNK to tune in the presence of HLA-A3/A11 on maternal cells was not taken into account in the assessments made, since the data on the inability of KIR3DL2 to tune49 were obtained on NK cells from peripheral blood and the applicability of these data to dNK is not obvious, in particular due to the distinct expression of KIR on dNK64.

### Modeling the effect of random iKIR genes expression on the representation of dNK functional phenotypes

Based on the presence of *N*_iKIR_ гe_HOB_ iKIR iKIR genes in the mother, their expression was described by an *N*_iKIR_-dimensional vector 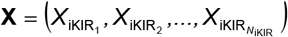of random variables 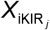, each of which has a Bernoulli distribution

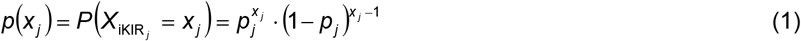

with binary outcomes *x* _*j*_ ∈ {0,1} (0 - no expression, 1 - expression). Vector **X** has 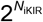 realizations 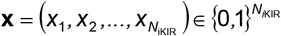,each of which corresponds to the expression of some set of the total number of maternal iKIR genes and some functional phenotype dNK_F_ (dNK_hypo_, dNK_HvG_, dNK_HvH_ или dNK_BD_). Probability density function 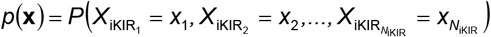 follows multivariate Bernoulli distribution

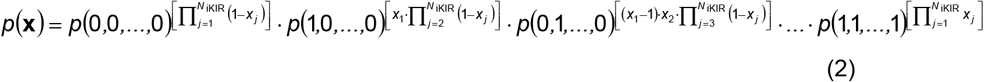

In general, the components of the vector **X** may be dependent random variables with different parameter *p*_*j*_ of the marginal Bernoulli distributions (1). But for a theoretical assessment of the possible influence of iKIR gene expression intensity on the representation of dNK_F_ populations, it is advisable to assume that the components of vector **X** are independent and identically distributed random variables with a Bernoulli distribution (*p* _*j*_ = *p* for any *j*). Then expressions (1) and (2) respectively take the form

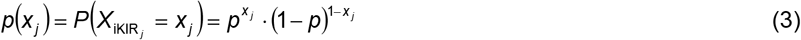

and

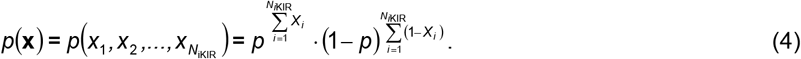

Determination of dNK functional phenotypes for all possible combinations of maternal iKIR genes in each pregnancy (see the section “Description of alloimmune response involving dNK” in Methods) allows us to assign to each **x** the value of the functions of the presence of functional phenotypes *dNK*_F_(**x**) (*dNK*_hypo_(**x**), *dNK*_HvG_(**x**), *dNK*_HvH_(**x**) и *dNK*_BD_(**x**)). Moreover, *dNK*_F_(**x**)=1 only for the assigned phenotype and *dNK*_F_(**x**)=0 for all others. The proportion of dNKs having a certain functional phenotype dNK_F_ during pregnancy is equal to the mathematical expectation *E*_F_ (*p*) of acquiring this functional phenotype, which is a function of *p*

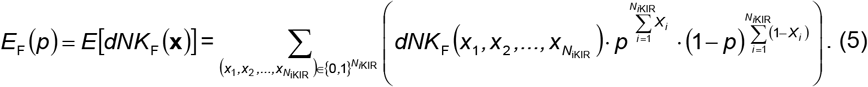

For presentation as percentages, the values *E*_F_ (*p*) calculated using equation (5) were multiplied by 100. Examples of theoretical dependencies of *E*_hypo_ (*p*), *E*_HvG_ (*p*), *E*_HvH_ (*p*) и *E*_BD_ (*p*) on the expression probability *p* of iKIRs for individual pregnancies with different *N*_iKIR_ and different numbers of affine pairs of iKIR and iKIR-L are shown in Supplementary Fig.3. For clinical observations from one group, including *N* pregnancies, the mean expectation value 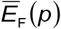 of the proportion of dNK with the functional phenotype dNK_F_ was calculated

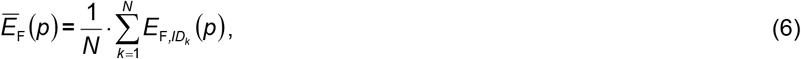

where 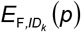 is the value of *E*_F_ (*p*) for the patient with the identifier *ID*_*k*_.

The assumption that the components of the vector **X** are independent and identically distributed Bernoulli random variables (*p* _*j*_ = *p* for any *j*) results in the possibility of describing the amount of iKIRs simultaneously acquired by dNK as a random variable *X* with a binomial distribution *X ∼ B*(*N*_iKIR_, *p*) and probability function

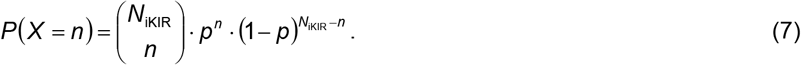

All model calculations, the results of which are presented in this work, were made using Microsoft Office Excel 2016 (Microsoft, Redmond, WA, USA).

### Statistical analysis

Statistical analysis was performed using Microsoft Office Excel 2016 (Microsoft, Redmond, WA, USA), MedCalc^®^ 14.8.1 (MedCalc, Ostend, Belgium), and Prism 10.2.3 (GraphPad, Boston, MA, USA). For statistical evaluation of differences in quantitative parameters, nonparametric Kruskal-Wallis and Friedman tests with Dunn’s post hoc test for paired comparisons were used. Comparison of the frequency of signs in clinical groups was made according to the Pearson’s χ2 goodness-of-fit test using Fisher’s exact test for post-hoc testing in paired comparisons. Null hypotheses were rejected and differences were considered significant at *P* < 0.05. For paired comparisons in post hoc analysis, differences were considered significant taking into account the Bonferroni correction.

Clustering was performed using WOLFRAM MATHEMATICA 13.0 (Wolfram, Champaign, IL, USA). The t-distributed stochastic neighbor embedding (t-SNE) method was used for dimensionality reduction. Clustering was performed using the k-medoids algorithm. The statistical significance of differences in the distribution of patients across clusters was assessed by analyzing contingency tables and the contingency coefficient (*C*_C_) using Pearson’s χ^2^ test.

## Acknowledgements

We thank all members of the Laboratory of Clinical Immunology, Laboratory of Bioinformatics, and Laboratory of Human Histocompatibility Genetics for critical reading of the manuscript and for providing feedback.

This research received no external funding.

## Ethics declarations

### Institutional Review Board Statement

The study was conducted in accordance with the Declaration of Helsinki, and approved by the Institutional Ethics Committee of National Medical Research Center for Obstetrics, Gynecology and Perinatology, named after academician V.I. Kulakov of the Ministry of Healthcare of the Russian Federation (protocol #35, dated 9 April 2009).

### Informed Consent Statement

Informed consent was obtained from all subjects involved in the study.

## Contributions

S.P.K. and N.K.T. conceived the idea for this article.

S.P.K., N.K.T. and L.V.K. designed, analyzed and interpreted experiments, as well as drafted the manuscript.

O.V.K., T.E.Y., Y.V.S. and P.I.B. performed experiments and/or analyzed and helped interpret data.

D.Y.T. aided in discussions of the project, data and manuscript.

G.T.S. supervised the project.

---

**Extended Data Fig. 1.**
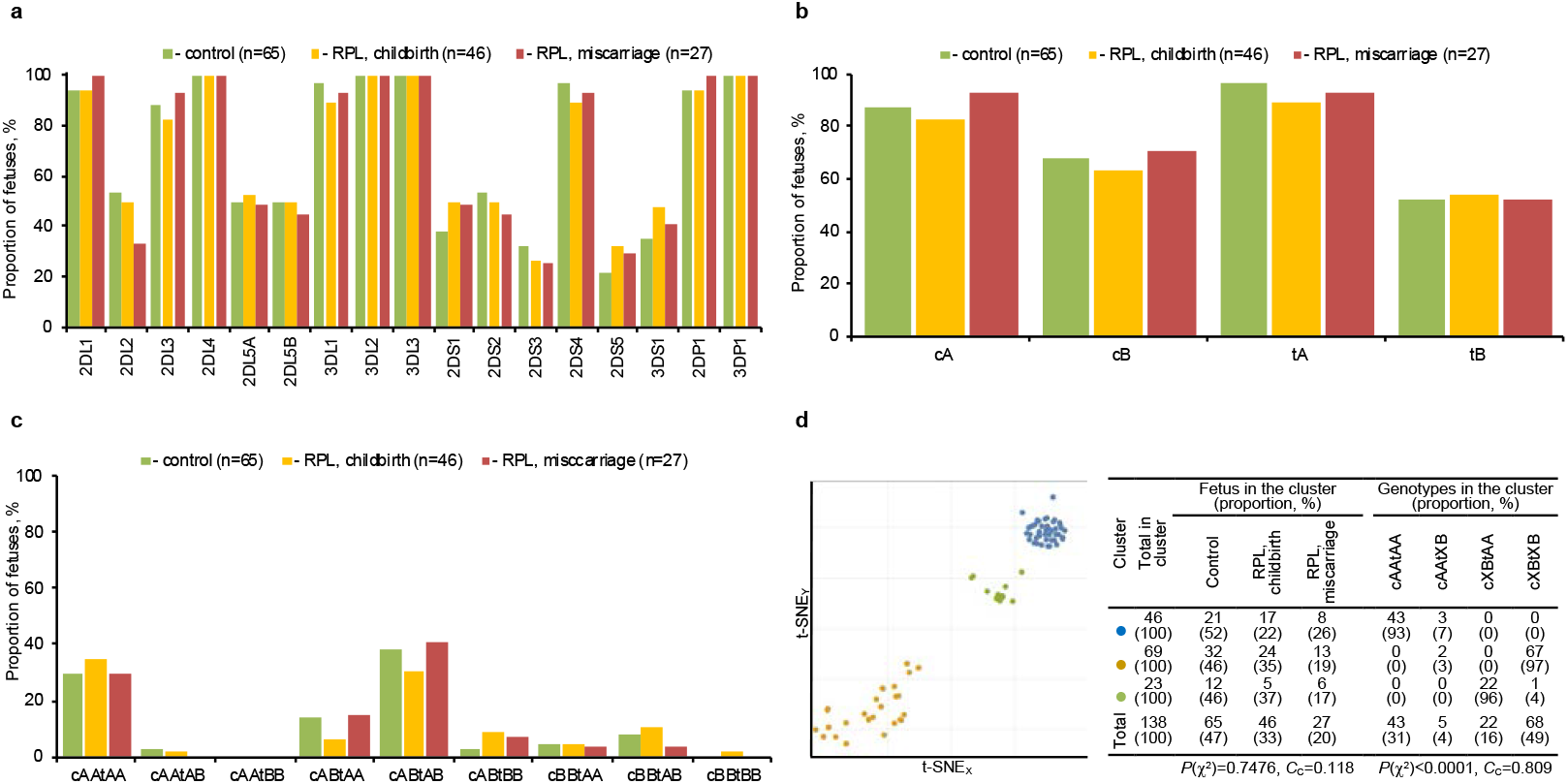
Fetal KIR genes and RPL. **a**, Proportion of fetuses with KIR genes in clinical groups. According to the χ^2^ test (*P*(χ^2^) > 0.05), there are no statistically significant differences between the clinical groups for all genes. **b**, Proportion of fetuses with combinations of KIR genes included in half-haplotypes A and B according to Fiq.1c. **c**, Proportion of fetuses with KIR genotypes determined by half-haplotypes identified in **b**. There are no statistically significant differences between the clinical groups according to the χ^2^ test (*P*(χ^2^) > 0.05) for all half-haplotypes (**b**) and all genotypes (**c**). **d**, Clustering of fetuses in the space of KIR genes (left diagram) and distribution of patients and genotypes in clusters (right table). The percentage of the total number of patients in the corresponding cluster is shown in parentheses in the cells of the table. The use of XB in the designation of genotypes through half-haplotypes means the summation of genotypes with the corresponding half-haplotypes AB and BB. The results of the χ^2^ tests are shown under the table data on the clusters distribution of patients from clinical groups and patients with genotypes.

**Extended Data Fig. 2.**
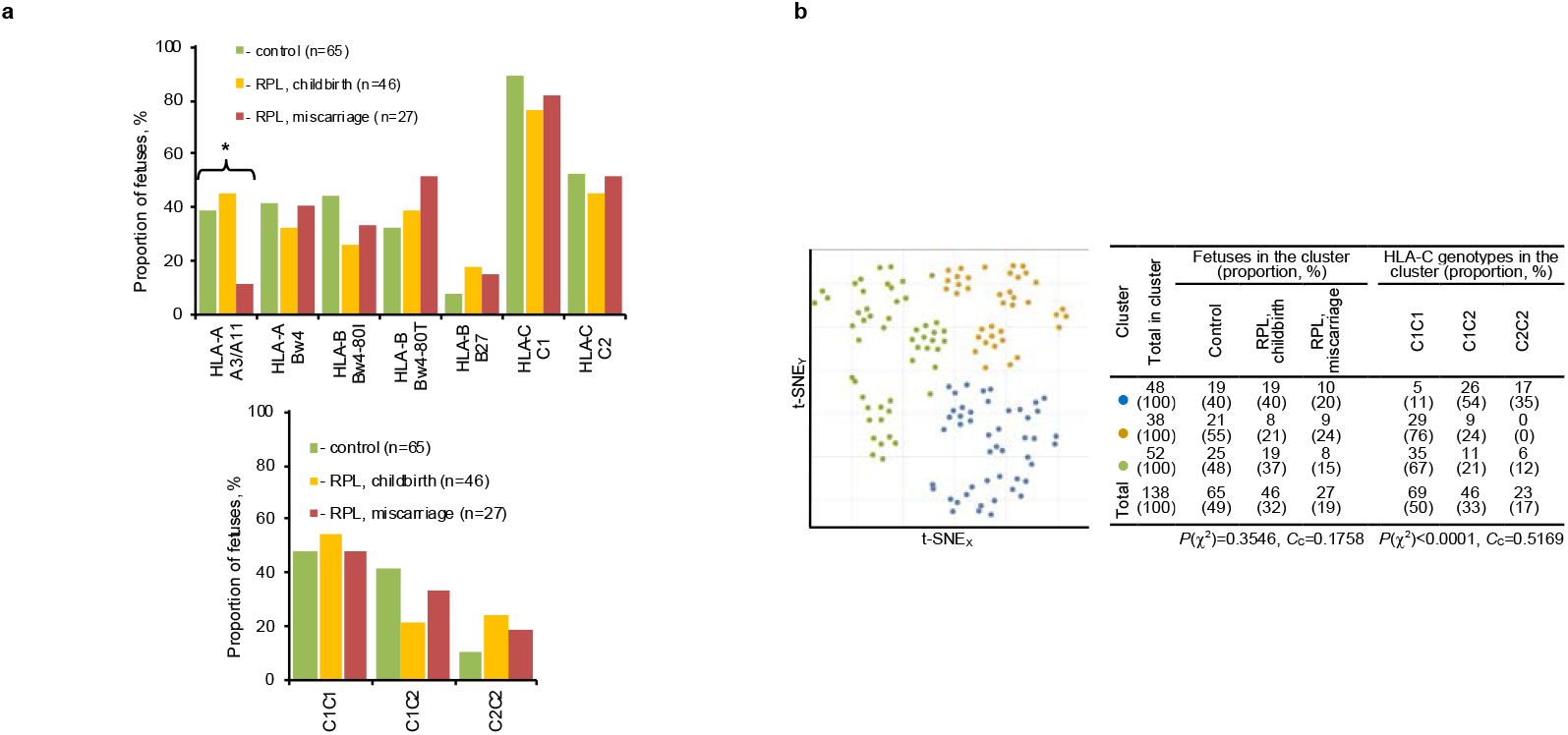
Fetal KIR-L and RPL. **a**, Proportion of fetuses with KIR-Ls (left diagram) and HLA-C genotypes (right diagram). According to the χ^2^ test, there are no statistically significant differences between the clinical groups for all KIR-L and HLA-C genotypes except HLA-A3/A11. * - *P*(χ^2^) = 0.0094; Fisher’s exact test for control versus RPL, miscarriage – *P* = 0.0120; Fisher’s exact test for RPL, childbirth versus RPL, miscarriage – *P* = 0.0039. Fisher’s exact test is significant with Bonferroni correction. **b**, Clustering of fetuses in the space of their own KIR-Ls (left diagram) and distribution of patients and HLA-C genotypes in clusters (right table). The percentage of the total number of patients in the corresponding cluster is shown in parentheses in the cells of the table. The results of the χ^2^ tests are shown under the table data on the clusters distribution of patients from clinical groups and patients with genotypes.

## Supplementary information

**Supplementary Table 1.** Clinical and anamnestic characteristics of mothers.

Clinical and anamnestic characteristics of mothers
| Characteristic | Control | RPL |  |
| --- | --- | --- | --- |
|  |  | childbirth | miscarriage |
| Number of mothers in the group | 65 | 44 | 23 |
| Age (years, median (Q1;Q3)) | 30 (27;34) | 33.5 (30;35) | 31 (28.8;34.3) |
| Diseases of the circulatory system<sup>a</sup> | 1 (1.5%) | 1 (2.3%) | 5 (21.7%)* |
| Endocrine disease<sup>a</sup> | 4 (6.2%) | 0(0%) | 1 (4.3%) |
| Diseases of the visual system<sup>a</sup> | 14 (21.5%) | 5 (11.4%) | 16 (69.6%)* |
| Diseases of the digestive system<sup>a,b</sup> | 14 (21.5%) | 1 (2.3%)* | 5 (21.7%) |
| Diseases of the urinary system<sup>a,b</sup> | 12 (18.5%) | 4 (9.1%) | 14 (60.9%)* |
| Fibromyomas disorder<sup>a,c</sup> | 7 (10.8%) | 2 (4.5%) | 5 (21.7%) |
| Endometriosis<sup>a,c</sup> | 4 (6.2%) | 1 (2.3%) | 2 (8.7%) |
| Female pelvic inflammatory diseases<sup>a,d</sup> | 5 (7.7%)* | 18 (40.1%) | 12 (52.2%) |
| Sexually transmitted diseases<sup>a,d</sup> | 14 (21.5%) | 15 (34.1%) | 4 (17.4%) |
| Total number of pregnancies before inclusion in the study | 65 | 182 | 106 |
| Induced abortion before inclusion in the study<sup>e</sup> | 0(0%)** | 22 (12.1%) | 7 (6.6%) |
| Total number of miscarriages up to 21 weeks 6 days before inclusion in the study<sup>e</sup> | 0(0%)* | 160 (87.9%) | 99 (93.4%) |

Q1;Q3, the 25th percentiles and 75th percentiles, respectively.
\*, $P < 0.05$ when compared between this group and each of the other groups. Continuous or discrete data compared using nonparametric Kruskal-Wallis test with Dunn's post-hoc test. Comparison of the proportions was performed using the Pearson's Chi-squared test and Fisher's exact test for post-hoc testing. $P$ values were corrected using the Bonferroni method.
\*\*, $P < 0.05$ when compared this group and RPL with childbirth group. Comparison of the proportions was performed as described above under this table.
<sup>a</sup>Number of mothers with diagnosis. Percentages are calculated as a proportion of the number of patients in the group.
<sup>b</sup>Mothers with current non-infectious and non-inflammatory diseases and mothers with cases of infectious and inflammatory diseases reflected in the medical history and cured before inclusion in the study.
<sup>c</sup>In mothers included in the study, the extent of the lesion did not contraindicate pregnancy according to clinical guidelines.
<sup>d</sup>Mothers with cases of infectious and inflammatory diseases reflected in the medical history and cured before inclusion in the study.
<sup>e</sup>Percentages are calculated as a proportion of the total number of pregnancies in the group.

**Supplementary Table 2.**
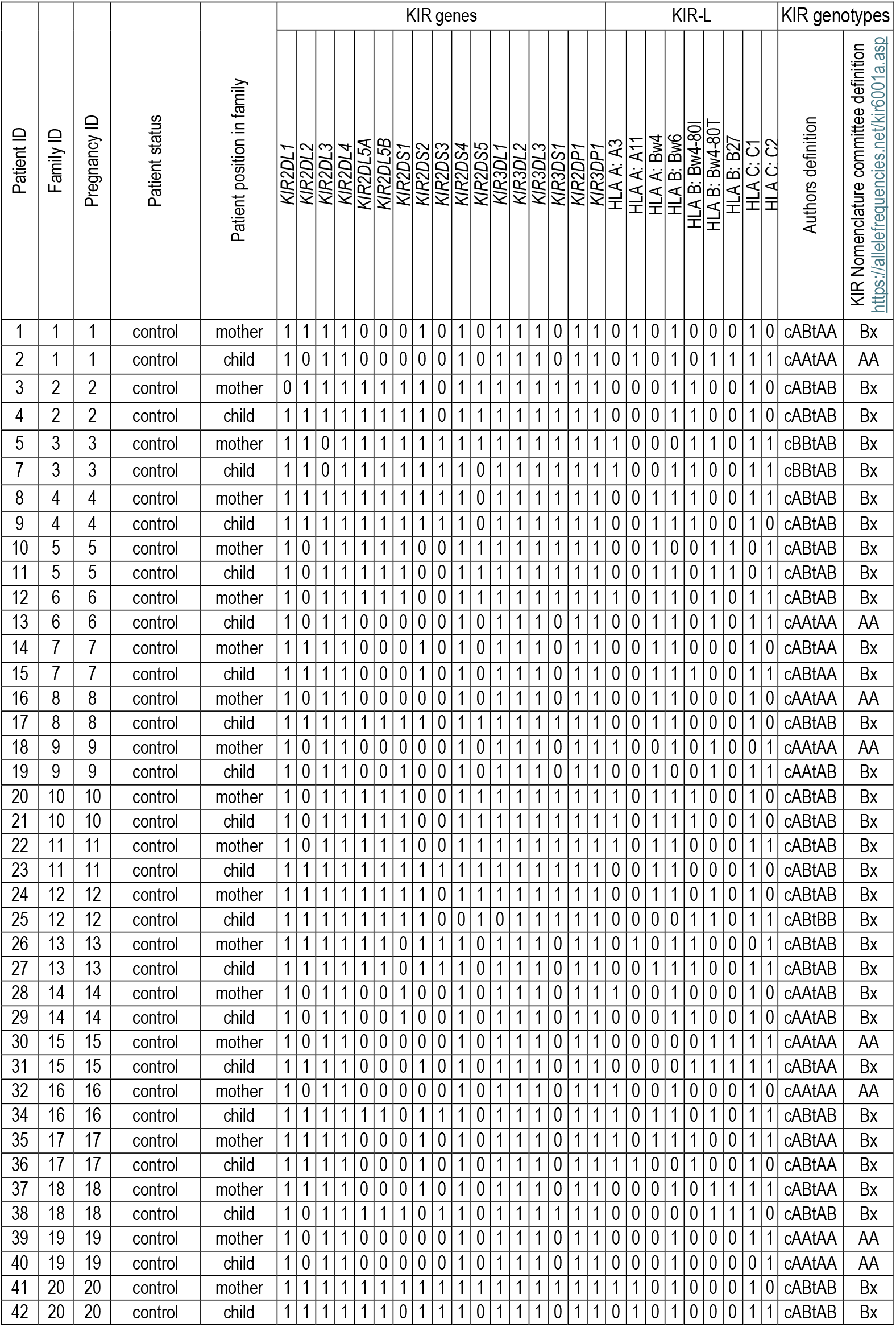
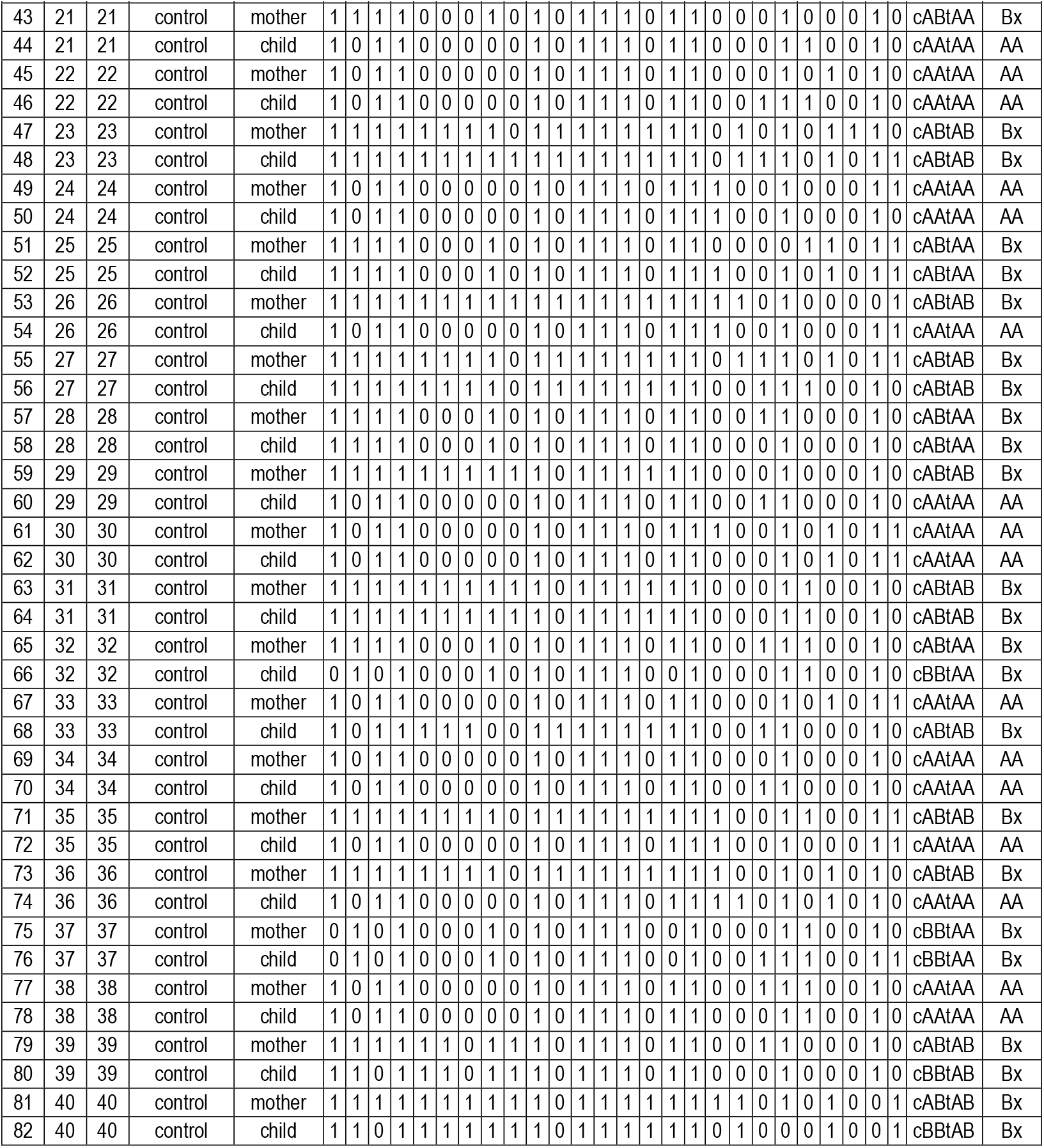
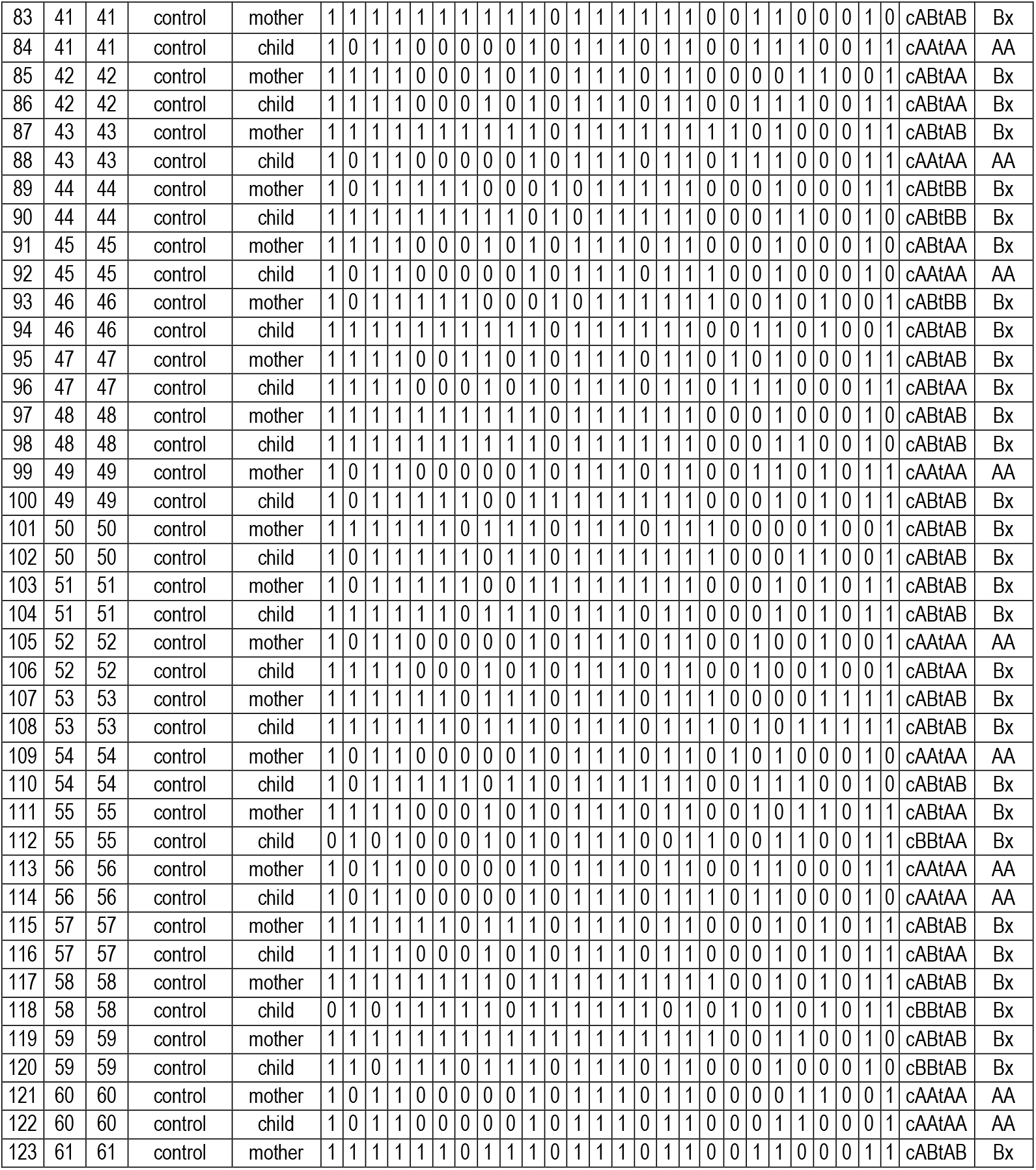
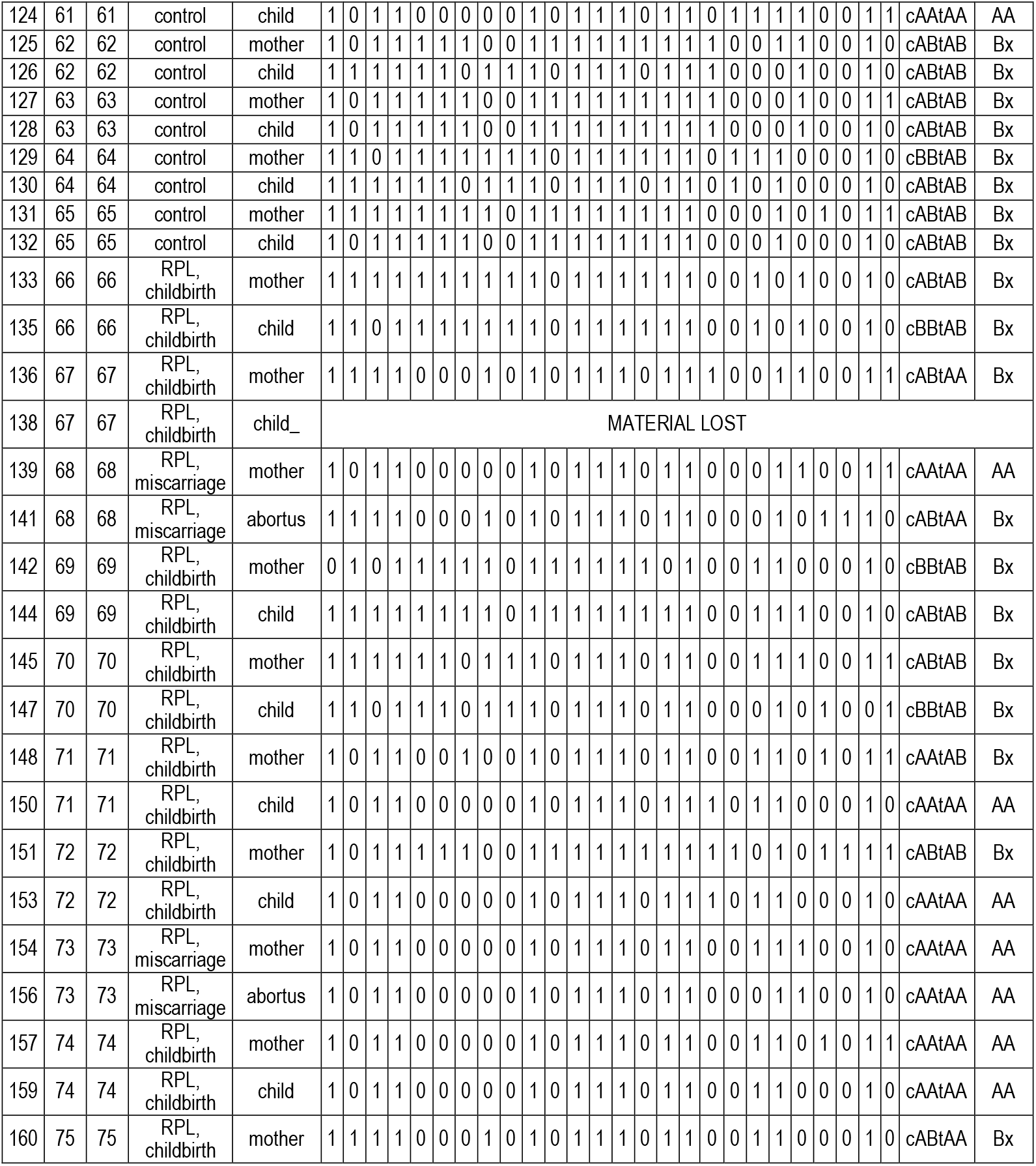
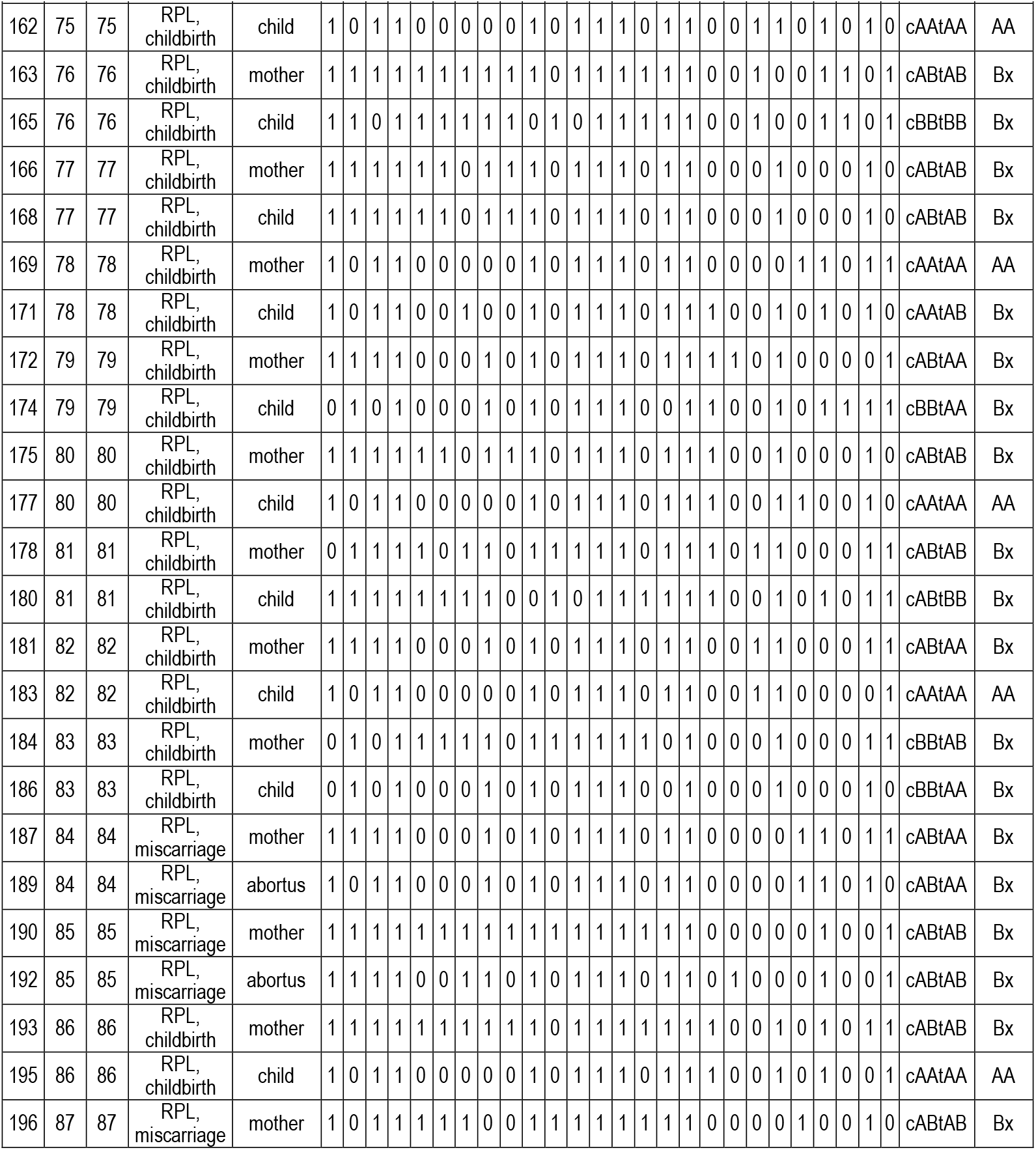
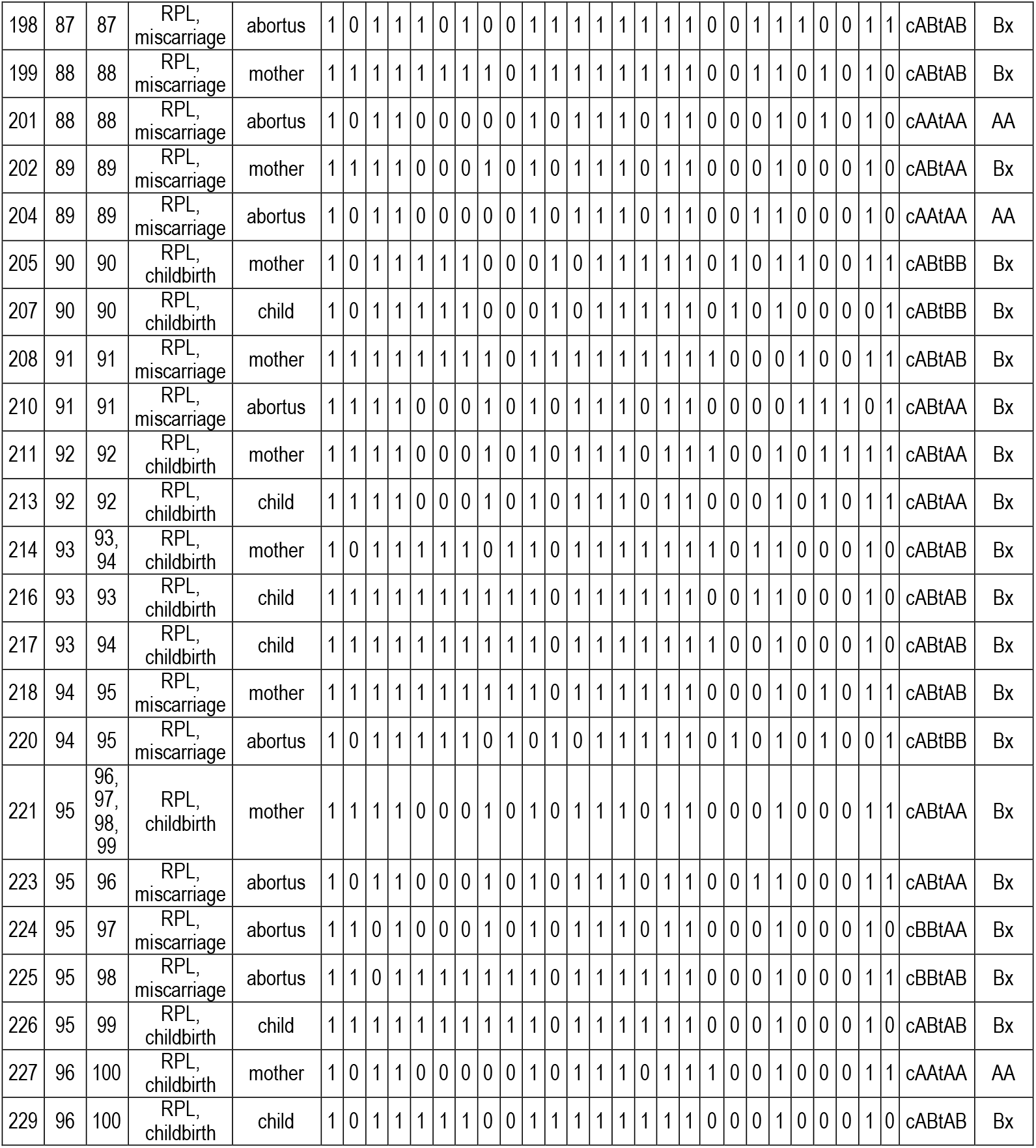
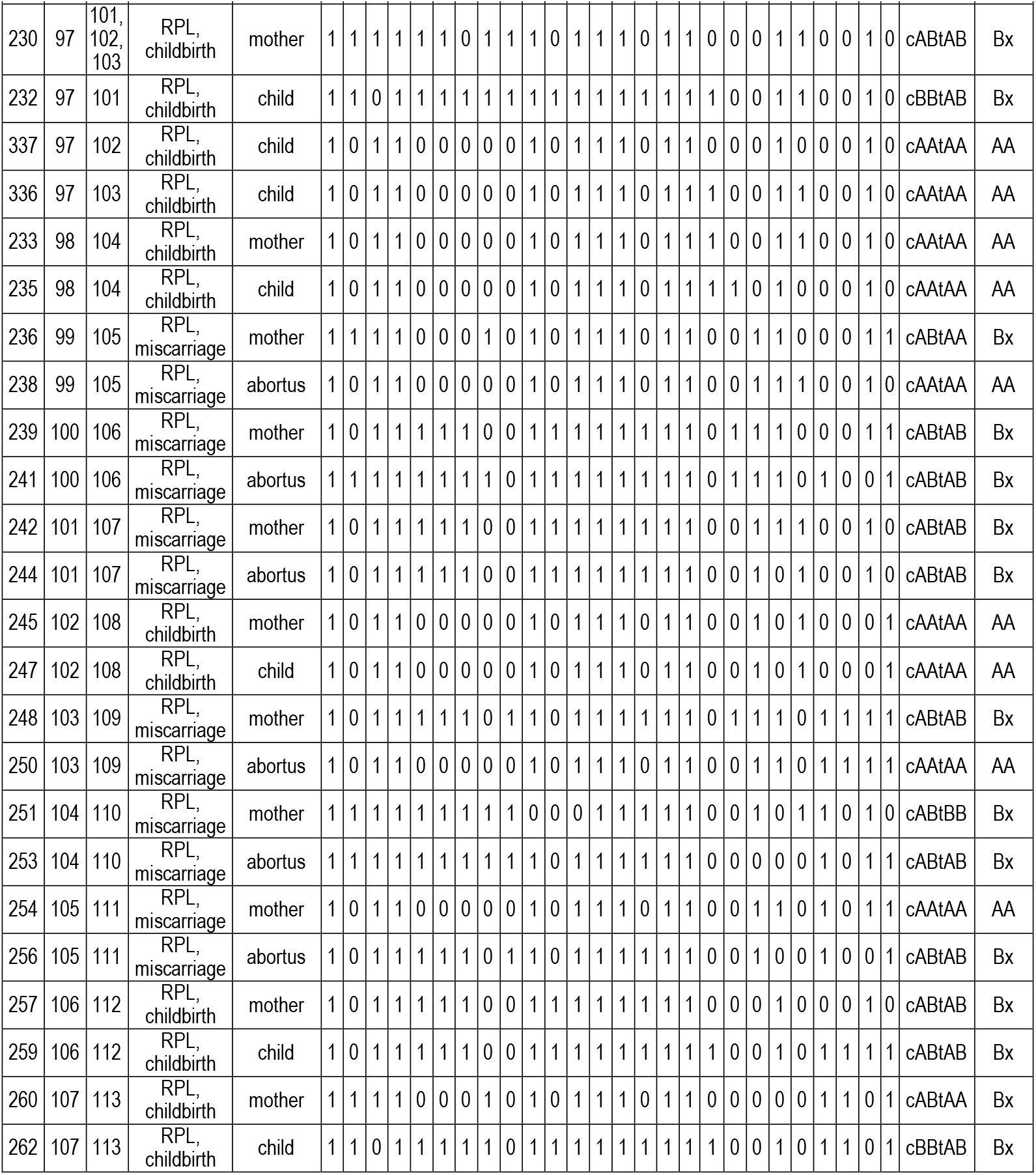
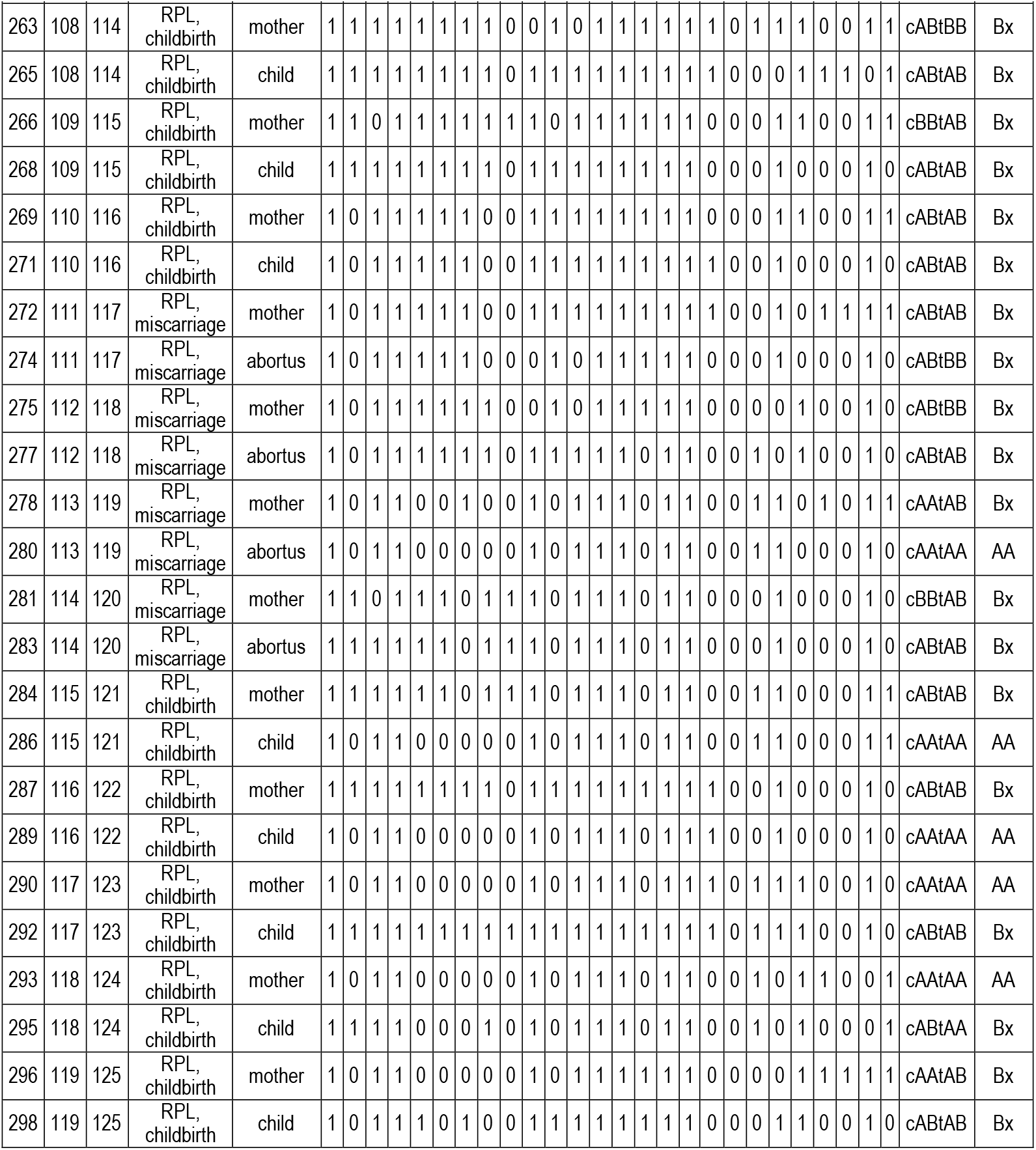
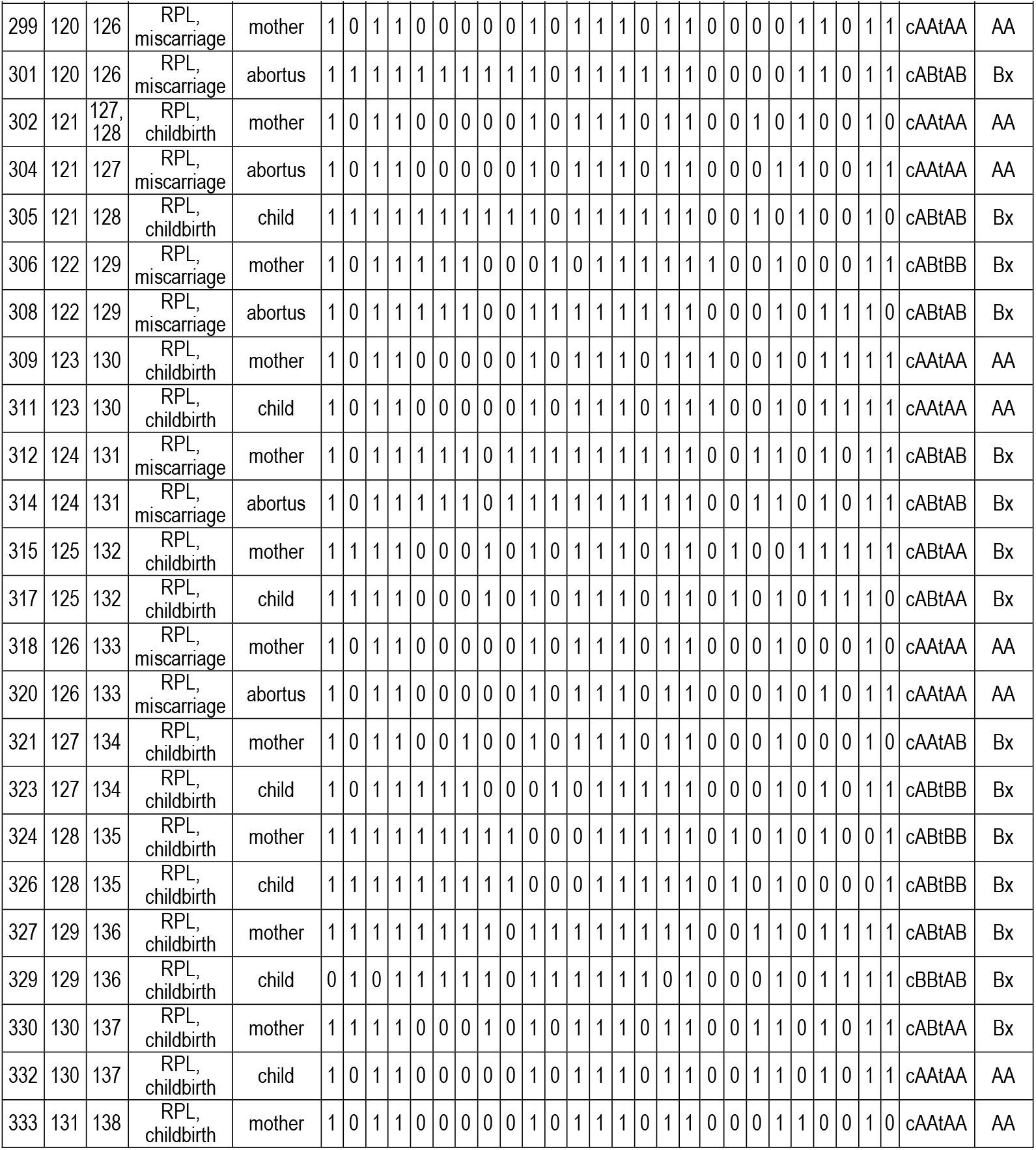
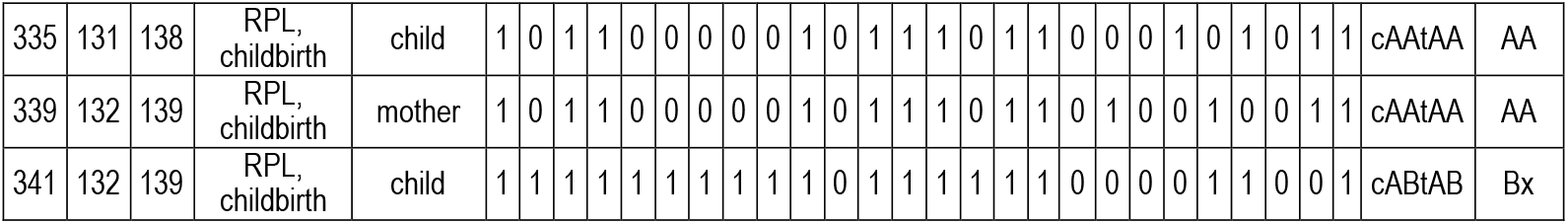
KIR genes and KIR-L of mothers and fetuses.

**Supplementary Fig. 1.**
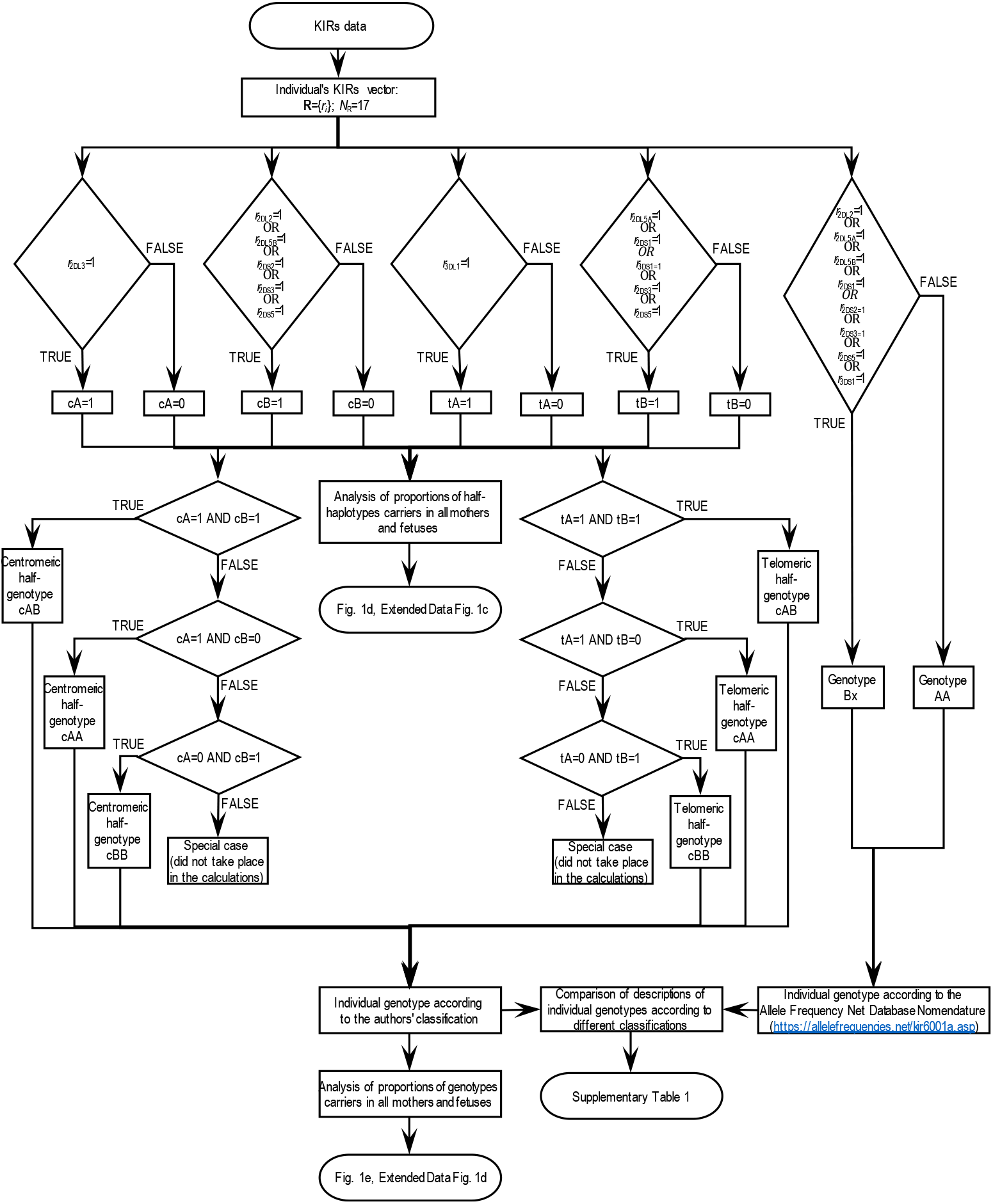
Algorithm for determining the presence of centromeric and telomeric A and B half-haplotypes of KIR and determining the genotypes of KIR

**Supplementary Fig. 2.**
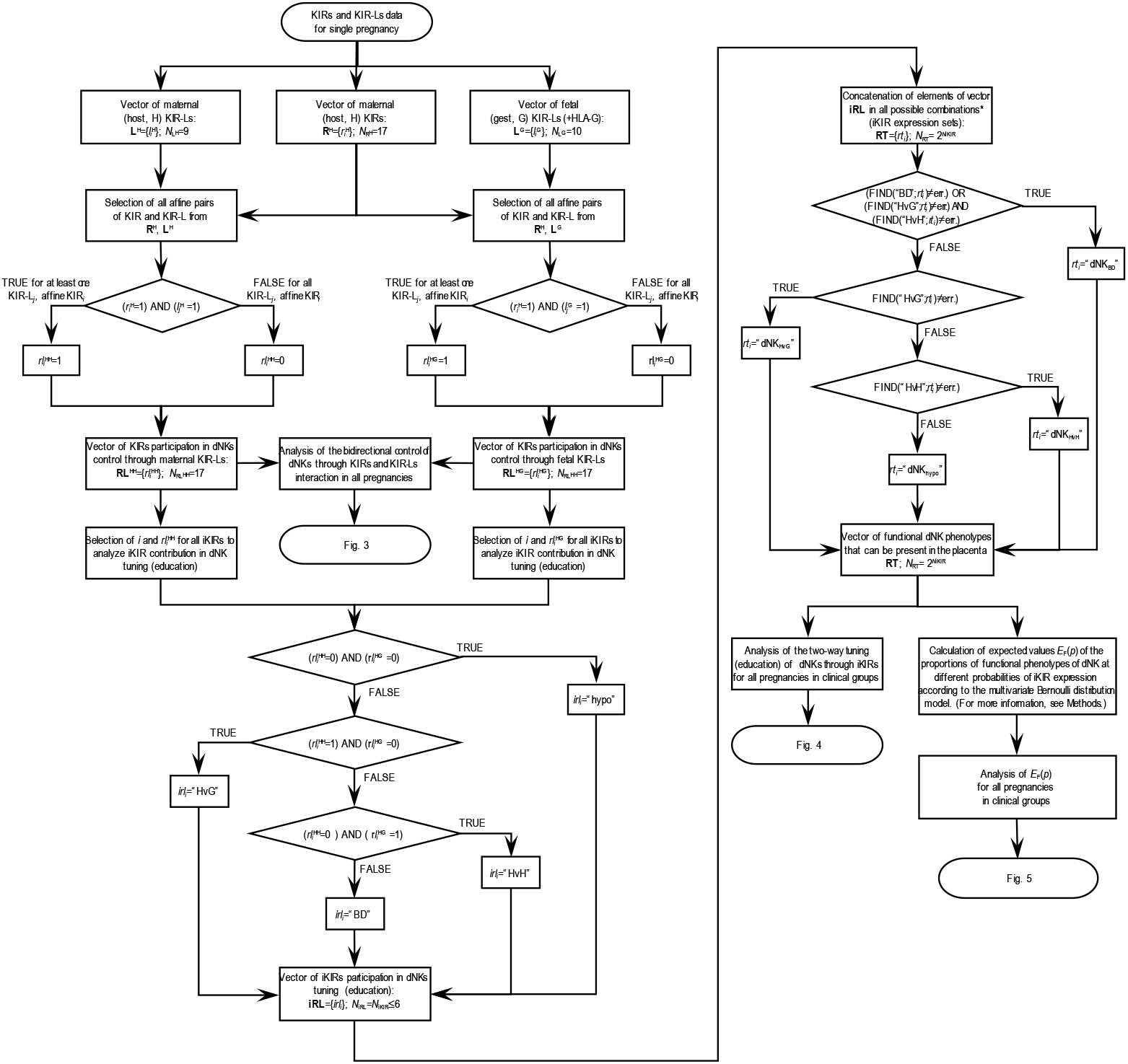
Algorithm for the approach to the description of the alloimmune response involving dNK. * - *rt*_*i*_ are text variables consisting of space-separated symbols of functional phenotypes for iKIR genes that correspond to 1 in the multivariate random variable 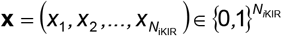 (e.g. “HvH”, “HvH hypo HvG”, “GvG BD BD GvH BD hypo”).

**Supplementary Fig. 3.**
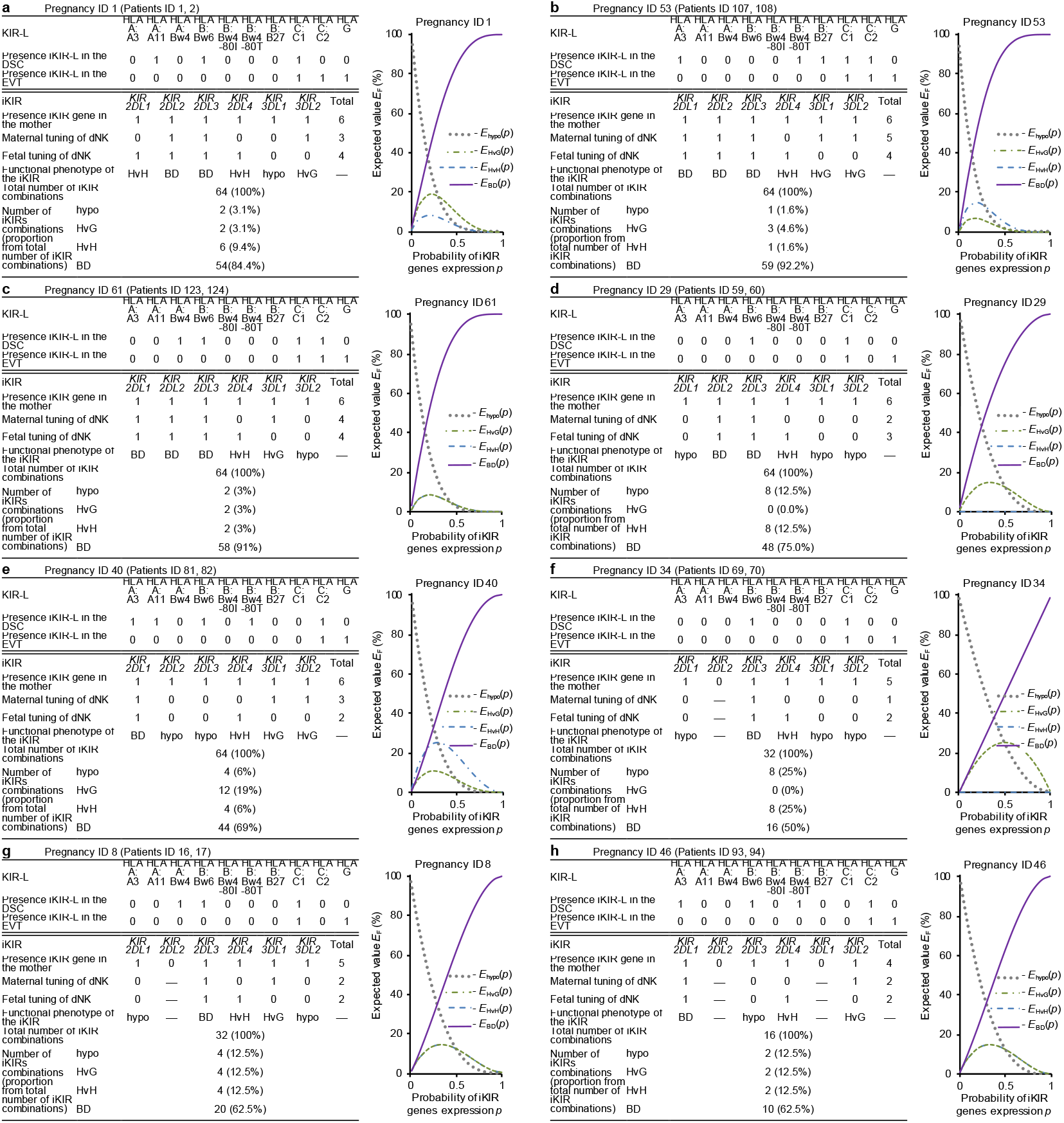
Examples of functional phenotypes estimation and hypothetical expected value *E*_F_ (*p*) of the proportions of dNK functional phenotypes at different probabilities *p* of iKIR genes expression for individual physiological pregnancies. **a**-**h**, For the iKIR and iKIR-L genes, one indicates presence and zero indicates absence. For dNK tuning using iKIR, one means tuning in the presence of affine iKIR and iKIR-L, zero means no tuning in the presence of iKIR due to the absence of iKIR-L, and a dash means no iKIR. Functional phenotype determination for individual iKIR gene and gene combinations, and hypothetical expected value *E*_F_ (*p*) calculations are described in Methods. **a**-**e**, cases with 6 maternal iKIR, **f**-**g**, cases with 5 maternal iKIR, and **h**, case with 4 maternal iKIR.

**Supplementary Fig. 4.**
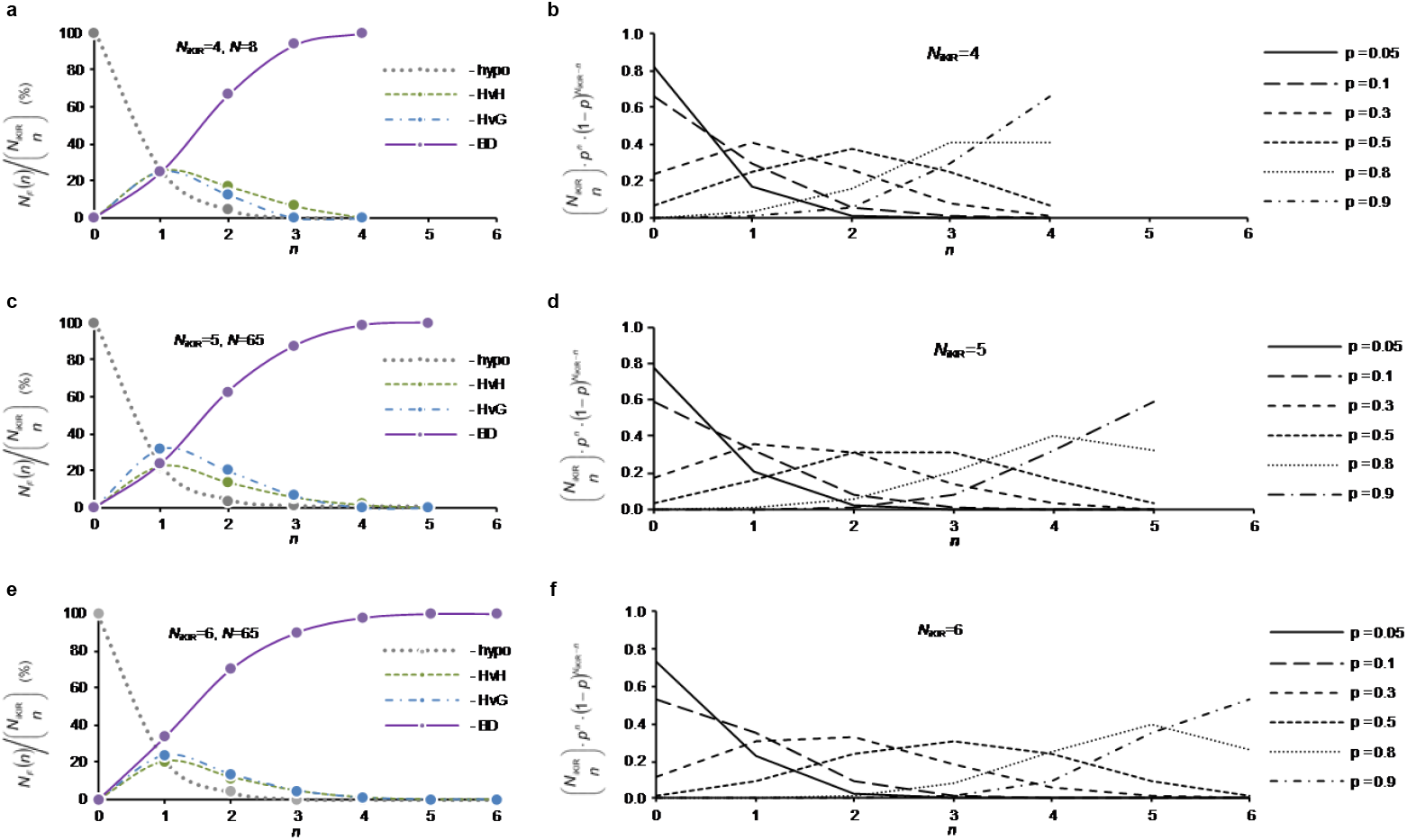
Functional phenotypes of dNKs at different numbers of expressed iKIRs and the use of the binomial distribution to describe the effect of expression probability on the distribution of the number of iKIRs acquired by dNK. **a, c** and **e**, average of proportion of number *N*_*F*_ (*n*) of iKIR combinations with different functional phenotypes from the total number 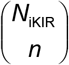 of possible iKIR combinations when expressing n of the *N*_iKIR_ genes for *N*_iKIR_=4 (**a**), *N*_iKIR_=5 (**c**) и *N*_iKIR_=6 (**e**). The calculations were done for all pregnancies without taking into account belonging to clinical groups. *N* is the number of pregnancies. **b, d** and **f**, binomial probability functions 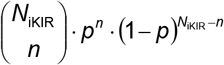 with different *p* describing the probability of dNK acquiring *n* iKIRs from *N*_iKIR_ for *N*_iKIR_=4 (**b**), *N*_iKIR_=5 (**d**) и *N*_iKIR_=6 (**f**). This model is based on the assumption that iKIR genes are expressed independently with equal probability *p* (see Methods for details).

**Supplementary Fig. 5.**
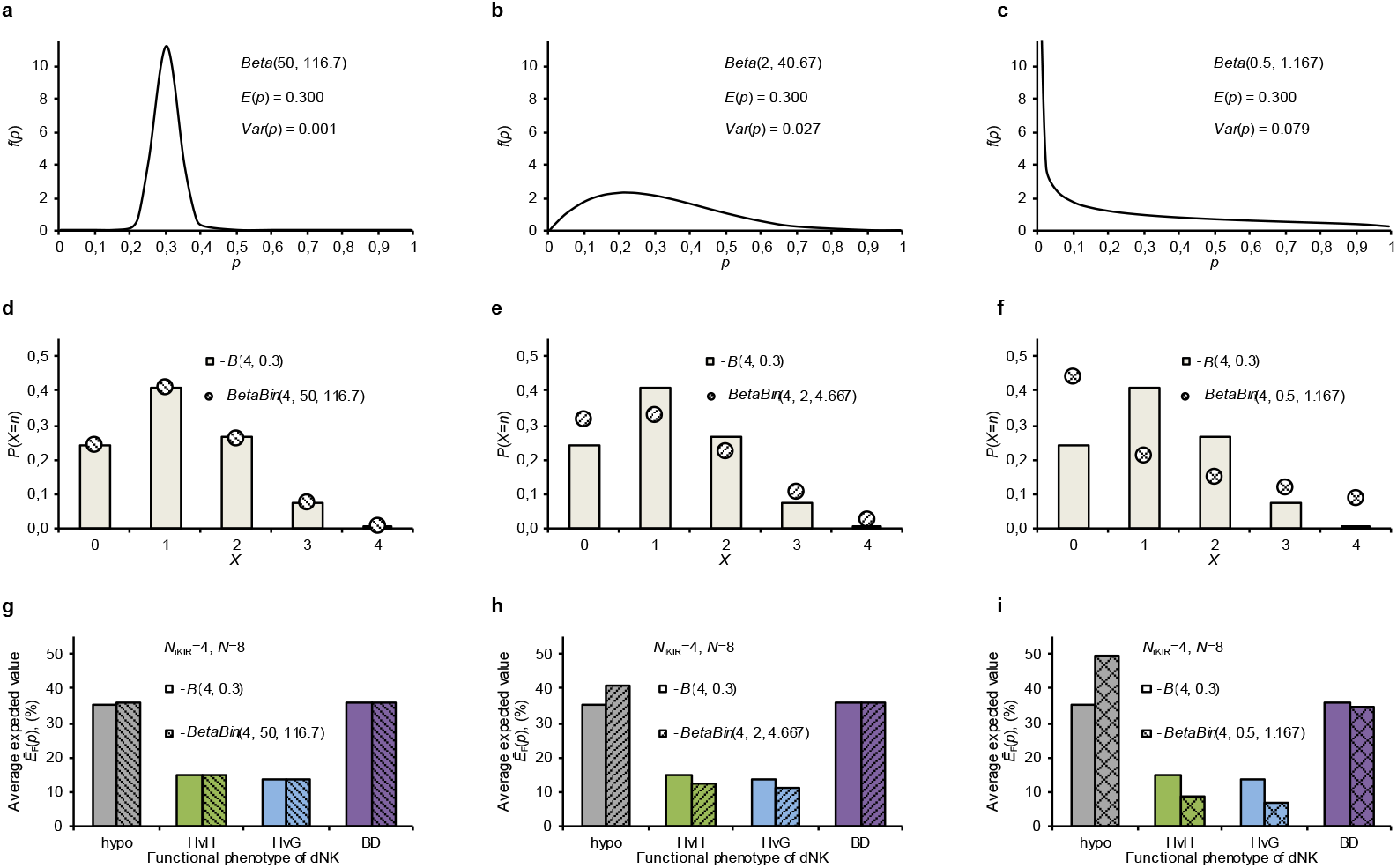
The variation in the probability of iKIRs expression between dNKs may influence the distribution of the amounts of iKIRs acquired by dNKs and lead a deviation from the product rule when iKIRs are independently expressed in an individual dNK. The probability distribution of *p* ∈ [0,1] of iKIRs expression among dNKs is modeled by a beta distribution *Beta*(*α, β*) with a probability density function 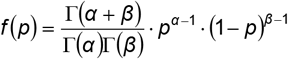, where 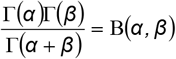 is the beta function and Γ(*z*) is the gamma function1. The compound beta-binomial distribution for describing the probability of expression of n from *N*_iKIR_ iKIRs has a probability function 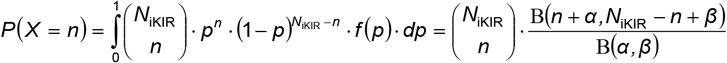. In the modeling, the choice of *N*_iKIR_=4 due to the presence of this number of iKIRs in publications with quantitative estimates of the probability of their expression3,4,5, and 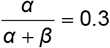 is close to the values of *p* at the maxima of the functions 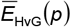 and 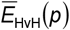 in the diagrams in Fig.5b. The diagrams in this Supplementary Fig. 5, placed one below the other, use the same *α* values and the same *β* values. **a, b** and **c**, examples of beta distributions with the same means (expectation values) 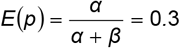 but different variances 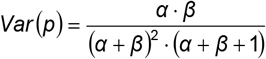, increasing from **a** to **b** and then to **c**. **d, e** and **f**, demonstration of a deviation from the product rule (deviation from the binomial distribution) with independent iKIRs expression and with a spreading in the iKIR expression probability between dNKs (beta-binomial distribution). The binomial distribution is shown by columns, and the beta-binomial distribution by circles. An increase in the *Var* (*p*) value, corresponding to an increase in the variability in the iKIRs expression probability between dNKs, is accompanied by a more pronounced violation of the product rule from **d** to **e** and then to **f**. **g, h** and **i**, estimating the possible effect of variability in the probability of iKIRs expression between dNKs on the mean expectation value 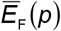 of the proportion of dNKs with functional phenotype F (hypo, HvG, HvH, BD) based on experimental data for pregnancies with maternal *N*_iKIR_=4. To estimate 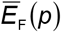, as in Supplementary Fig. 4a, all cases of pregnancies with maternal *N*_iKIR_=4 were used, without taking into account their membership in clinical groups. The calculation of 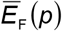 is described in Methods. An increase from **g** to **h** and then to **i** in the spread of the probability of iKIRs expression between dNKs (an increase in the value of *Var* (*p*)) leads to smaller proportions of dNK_HvH_ and dNK_HvG_ actively participating in the formation of the placenta than with a binomial distribution corresponding to *Var* (*p*) = 0, and this shift in proportions increases with increasing *Var* (*p*). The negative skew of the beta-binomial distribution used to model the variation in *p* at 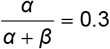 causes, simultaneously with a decrease in the proportions of dNK_HvH_ and dNK_HvG_, an increase in the proportion of hyporeactive dNK_hypo_ with a slight decrease in “anti-infectious” dNK_BD_.

## Supplementary Note 1

### A model estimation of the possible effect of the HLA A3/A11 absence in the donor (guest) on the representation of functional phenotypes of NK cells of the recipient (host)

Table I shows model estimates of the representation of recipient (host) NK cells with functional phenotypes (hypo, HvG, HvH, BD) in two-way tuning by recipient and donor (guest) target cells according to the algorithm described in the Methods. The estimates were performed for the general case of recipient NK cells possessing KIR3DL2 (required due to the framework *KIR3DL2* gene), _S_iKIR (iKIR affine for _S_iKIR-L, present in the recipient) and _NS_iKIR (iKIR affine for _NS_iKIR-L, absent in the recipient). According to the modeling results, under otherwise equal conditions, the absence of HLA-A3/A11 in the donor leads to a higher proportion of iKIRs combinations that confer the HvG functional phenotype to the recipient NK cell and/or a lower proportion of iKIRs combinations that confer the HvH functional phenotype to the recipient NK cell.

**Table I.** Effect of HLA-A3/A11 in the donor (guest) on the presence of recipient (host) NK cells functional phenotypes in a model involving KIR3DL2 in the two-way tuning.

| Recipient (host) |  | Donor (gest)<br>iKIR-Ls<br>without<br>taking into<br>account<br>HLA A3/A11 | Proportion of iKIR combinations with functional phenotype<br>from the total number of all possible iKIR combinations (n=8)<br>for recipient NK cells, % |  |  |  |  |  |  |  |  |  |  |  |
| --- | --- | --- | --- | --- | --- | --- | --- | --- | --- | --- | --- | --- | --- | --- |
| iKIRs | iKIR-Ls |  | Donor (gest) iKIR-Ls<br>**with** HLA-A3/A11 |  |  |  | Donor (gest) iKIR-Ls<br>**without** HLA-A3/A11 |  |  |  | Donor (gest) iKIR-Ls<br>**without minus with**<br>HLA-A3/A11 |  |  |  |
|  |  |  | hypo | HvG | HvH | BD | hypo | HvG | HvH | BD | hypo | HvG | HvH | BD |
| KIR3DL2,<br>siKIR,<br>NSiKIR | HLA-A3/A11,<br>siKIR-L | siKIR-L,<br>NSiKIR-L | 12,5 | 0,0 | 12,5 | 75,0 | 12,5 | 12,5 | 12,5 | 62,5 | 0,0 | 12,5 | 0,0 | -12,5 |
|  |  | NSiKIR-L | 12,5 | 12,5 | 12,5 | 62,5 | 12,5 | 37,5 | 12,5 | 37,5 | 0,0 | 25,0 | 0,0 | -25,0 |
|  | siKIR-L | siKIR-L,<br>NSiKIR-L | 12,5 | 0,0 | 37,5 | 50,0 | 25,0 | 0,0 | 25,0 | 50,0 | 12,5 | 0,0 | -12,5 | 0,0 |
|  |  | NSiKIR-L | 12,5 | 12,5 | 37,5 | 37,5 | 25,0 | 25,0 | 25,0 | 25,0 | 12,5 | 12,5 | -12,5 | -12,5 |

$s_i$ KIR - iKIR of the recipient, affine for the $s_i$ KIR-L that recipient has; $ns_i$ KIR - iKIR of the recipient for which recipient does not have an affine $ns_i$ KIR-L. Exceeding of the predicted proportion of NK cells with a certain phenotype in the presence of HLA-A3/A11 in the donor (guest) cells over the proportion of NK cells with the same phenotype in the absence of HLA-A3/A11 in the donor cells is highlighted in pink. The reverse arrangement of these proportions is highlighted in blue.

Table II shows model estimates similar to those in Table I, assuming that KIR3DL2 is incapable of participating in NK cell tuning, but that this iKIR has an inhibitory effect with suppression of NK cell cytolytic activity against HLA-A3/A11 target cells1. In this case, it was only considered that NK cells with KIR3DL2, tuned through the interaction of any other iKIR with the affine iKIR-L, are not cytolytic against HLA-A3/A11-positive target cells. As a result, for some combinations of donor iKIR-Ls and recipient iKIR-Ls, higher HvG cytolytic activity of recipient NK cells against donor target cells is also predicted in the absence of HLA-A3/A11 in the donor.

**Table II.**
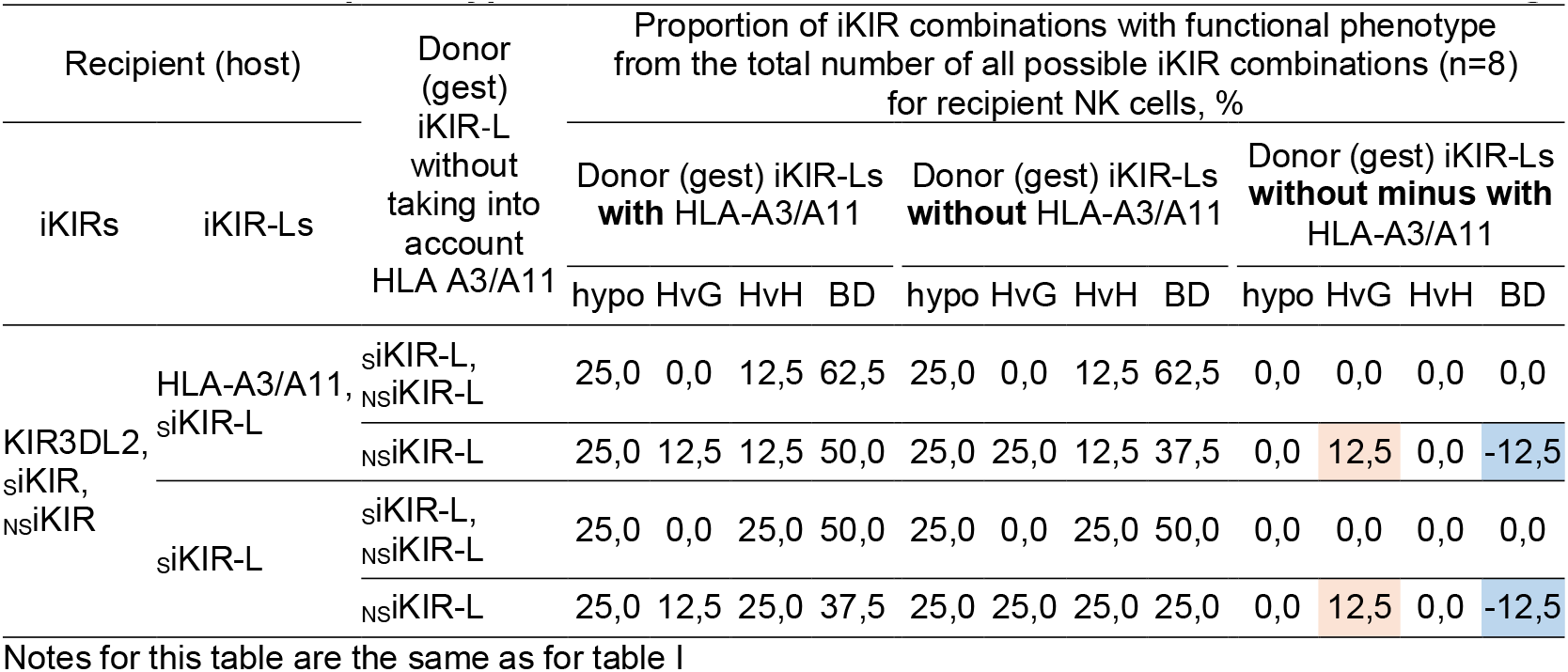
Effect of HLA-A3/A11 in the donor (guest) on the presence of recipient (host) NK cells functional phenotypes in a model without involvement of KIR3DL2 in the tuning.

Considering that the absence of HLA-A3/A11 in the donor corresponds to a higher probability of kidney transplant engraftment2, based on the data in Tables II and I, it can be assumed that the presence of KIR3DL2-related higher HvG alloreactivity or reduced HvH autoreactivity in host (recipient) NK cells under certain conditions may contribute to guest (graft) survival.

In addition to HLA-A3/A11, another affine iKIR-L, HLA B27, is known for KIR3DL23, the presence of which and its interaction with KIR3DL2 were taken into account in the analysis presented in the main text of the article. In the approach used by us, the presence of several affine iKIR-Ls for one iKIR suggests that such an iKIR is involved in NK cell tuning in the presence of any of the affine iKIR-Ls or their combinations. The absence of HLA-A and HLA-B expression on trophoblasts means the absence of HLA-A3/A11 and HLA-B27 expression on guest cells. Formally, this already corresponds to the conditions for better compatibility of the guest and the host, based on the data on renal engraftment2. However, according to Table I, in the absence of HLA-A3/A11 and HLA-B27 in the guest, the greatest number of combinations of iKIRs giving the NK cell a functional HvG phenotype involving KIR3DL2 is expected in the presence of HLA-A3/A11 and HLA-B27 in the host. In this regard, the assumption of better conditions for guest-host compatibility in the presence of HvG alloreactivity with the participation of KIR3DL2 allows us to expect better pregnancy gestation in the presence of HLA A3/A11 and HLA B27 in the mother, which is what took place in our study (Fig. 3a, Fig. 4e).

